# Mathematical modeling suggests cytotoxic T lymphocytes control growth of B16 tumor cells in collagin-fibrin gels by cytolytic and non-lytic mechanisms

**DOI:** 10.1101/2023.03.28.534600

**Authors:** Barun Majumder, Sadna Budhu, Vitaly V. Ganusov

## Abstract

Cytotoxic T lymphocytes (CTLs) are important in controlling some viral infections, and therapies involving transfer of large numbers of cancer-specific CTLs have been successfully used to treat several types of cancers in humans. While molecular mechanisms of how CTLs kill their targets are relatively well understood we still lack solid quantitative understanding of the kinetics and efficiency at which CTLs kill their targets in different conditions. Collagen-fibrin gel-based assays provide a tissue-like environment for the migration of CTLs, making them an attractive system to study the cytotoxicity in vitro. Budhu *et al.* [1] systematically varied the number of peptide (SIINFEKL)- pulsed B16 melanoma cells and SIINFEKL-specific CTLs (OT-1) and measured remaining targets at different times after target and CTL co-inoculation into collagen-fibrin gels. The authors proposed that their data were consistent with a simple model in which tumors grow exponentially and are killed by CTLs at a per capita rate proportional to the CTL density in the gel. By fitting several alternative mathematical models to these data we found that this simple “exponential-growth-mass-action-killing” model does not precisely fit the data. However, determining the best fit model proved difficult because the best performing model was dependent on the specific dataset chosen for the analysis. When considering all data that include biologically realistic CTL concentrations (*E* ≤ 10^7^ cell/ml) the model in which tumors grow exponentially and CTLs suppress tumor’s growth non-lytically and kill tumors according to the mass-action law (SiGMA model) fitted the data with best quality. Results of power analysis suggested that longer experiments (∼ 3 − 4 days) with 4 measurements of B16 tumor cell concentrations for a range of CTL concentrations would best allow to discriminate between alternative models. Taken together, our results suggest that interactions between tumors and CTLs in collagen-fibrin gels are more complex than a simple exponential-growth- mass-action killing model and provide support for the hypothesis that CTLs impact on tumors may go beyond direct cytotoxicity.

## Introduction

Cytotoxic T lymphocytes (**CTLs**) are important in controlling some viral infections and tumors [2, 3]. CTLs exhibit such control via several complimentary mechanisms among which direct cytotoxicity, the ability of CTLs to kill virus-infected or tumor (target) cells, is important. Killing of a target cell by a CTL in vivo is a multi-step process: 1) CTL must migrate to the site where the target is located, 2) CTL must recognize the target (typically by the T cell receptor (**TCR**) on the surface of T cells binding to the specific antigen presented on the surface of the target cell), 3) CTL must form a cytotoxic synapse with the target, and 4) CTL must induce apoptosis of the target cell by secreting effector molecules (e.g., perforin and granzymes) or through Fas/Fas-ligand interactions [4–8]. The relative contribution of these steps to the efficiency at which a population of CTLs kill their targets in vivo remains poorly understood especially in complex tissues. Improving efficacy of cancer based immunotherapies such as adoptive transfer of cancer-specific T cells will likely come from better understanding of a relative contribution of these processes to tumor control [9].

Many previous studies have provided quantitative insights into how CTLs eliminate their targets in vitro. First insights came from generating conjugates between target cells and CTLs and quantifying how quickly a target cell dies when either being bound by different number of CTLs, or when one CTL is bound to different targets [10–17]. Further in vitro studies highlighted that killing by CTLs may kill multiple targets rapidly [18–20] but also highlighted heterogeneity in efficacy at which individual CTLs kill their targets [21, 22]. Interestingly, killing of tumor cells in vitro may take long time (hours) with speed and turning being important in determining the likelihood that a CTL will find and kill the target [23, 24]. One study suggested that killing of targets in vitro may follow the law of mass-action [25]. Killing efficiency of CTLs has been also evaluated in so-called chromium release assays that have been a standard method in immunology to measure T cell cytotoxicity in vitro [26–33].

Evaluating killing efficacy of CTLs in vivo is challenging. One approach to evaluate how a population of CTLs eliminates targets in vivo has been to perform in vivo cytotoxicity assay [34]. In the assay two populations of cells, one pulsed with a specific peptide and another one being a control, are transferred into mice carrying peptide-specific CTLs, and the relative percent of peptide- pulsed targets is determined in a given tissue (typically spleen) after different times after target cell transfer [34–36]. Different mathematical models have been developed to determine specific terms describing how CTLs kill their targets and to estimate CTL killing efficacy; such estimates varied orders of magnitude between different studies often using similar or even same data [37–44]. One study suggested that mass-action killing term is fully consistent with data from different in vivo cytotoxicity experiments [42] while other studies based on theoretical arguments suggested that killing should saturate at high CTL or target cell densities [38, 45, 46].

Intravital imaging has provided additional insights into how CTLs kill their targets [47, 48]. One pioneering study followed interactions between peptide-pulsed B cells and peptide-specific CTLs in lymph nodes of mice and found that CTLs and their targets form stable conjugates and move together until the target stops and dies, presumably due to the lethal hit delivered by the CTL [49]. This and other studies revealed that to kill a target in vivo, CTLs either need to interact with the target for a long time or multiple CTLs must contact a target to ensure its death [3, 50–55]. Interestingly, killing of tumor cells or cells, infected with Plasmodium parasites, required hours that is longer than the killing time estimated from in vivo cytotoxicity assays [40, 51, 52, 55, 56]. This may be due to different levels of presented antigens (pulsed with a high concentration of a cognate peptide targets vs. targets expressing exogenous antigens) but may be also due to differences in intrinsic killing abilities of different T cells. Mathematical modeling provided quantification of how CTLs kill their targets and of various artifacts arising in intravital imaging experiments (e.g., zombie contracts) [57, 58]; we have recently suggested that killing efficacy of individual Plasmodium-specific CTLs is too low to rapidly eliminate a Plasmodium liver stage highlighting the importance of clusters of CTLs around the parasite for its efficient elimination [56].

Even though studying how CTLs kill their targets in vivo is ideal, such experiments are expensive, time-consuming, and low throughput. On the other hand, traditional in vitro experiments (e.g., on plates or in wells) suffer from the limitation that CTLs and targets do not efficiently migrate on flat surfaces as they do in vivo in many tissues. Collagen-fibrin gels have been proposed as a useful in vitro system to study CTL and target cell interactions that allows to better represent complex 3D environment of the tissues with low cost and higher throughput [1, 59, 60]. CTLs readily migrate in these gels with speeds similar to that of T cells in some tissues in vivo [61]. One recent study measured how CTLs, derived from transgenic mice whose TCRs are all specific for the peptide SIINFEKL (from chicken ovalbumin), can eliminate SIINFEKL peptide pulsed B16 tumor cells in collagen-fibrin gels [1]. Interestingly, the rate at which tumor cells were lost from the gel was linearly dependent on the concentration of CTLs in the gel (varied from 0 to 10^7^ cells/ml) and was independent of the number of B16 tumor cells deposited in the gel [1]. This result suggested that the killing of B16 tumor cells in collagen-fibrin gels follows the law of mass-action, and given that the population of B16 tumor cells grew exponentially with time, the authors proposed that 3.5 × 10^5^ cell/ml of CTLs are required to prevent B16 tumor cell accumulation in gels.

In this paper we more rigorously re-analyzed data published by Budhu *et al.* [1] along with two additional previously unpublished datasets on CTL killing of B16 tumor cells in collagen-fibrin gels. We found that the simple exponential growth and mass-action killing model never provided the best fit of the data, and which model (out of 4 tested) fitted the data best was dependent on the specific subset of the data used for the analysis. The model in which CTLs reduce the growth rate of B16 tumor cells and kill the tumors via a mass-action law (proportional to concentrations of the CTLs and tumors) fitted one largest dataset (431 gels) with best quality. Importantly, the type of the killing term was critical in predicting CTL concentration that would be needed to eliminate most of the tumor cells within a defined time period (100 days) suggesting the need for future experiments. Following our recent framework for experimental power analyses [62] we simulated various experimental designs and found that some designs would better allow to discriminate between alternative mathematical models of CTL-mediated control of B16 tumor cells, and thus, will allow to better predict how many CTLs are needed for tumor control.

## Materials and methods

### Experimental details and data

All main details of experimental design are provided in the previous publication [1]. In short, 10^3^*−*10^6^SIINFEKL-pulsed B16 melanoma tumor cells (=10^4^ *−* 10^7^ cell/ml) were inoculated alone or with 10^3^ *−* 10^6^ (equivalent to 10^4^ *−* 10^7^ cell/ml) of activated OT1 T cells (CTLs) into individual wells containing collagen-fibrin gels. At different times after co-inoculation of cells, gels were digested, and the resulting solution was diluted 10^1^ *−* 10^3^ fold (depending on the initial targeted B16 cell concentration) in growth medium, and the number of surviving B16 cells in each gel was counted [1]. The data are thus given as the concentration of B16 tumor cells (in cell/ml) surviving in the gels by a given time. Budhu *et al.* [1] provided us with the data from their published experiments (Datasets 1, 2, and 3) as well as two additional unpublished datasets (Datasets 4 and 5).

1. **Dataset 1** (growth): SIINFEKL-pulsed B16 melanoma cells were inoculated in a 3D collagen-I-fibrin gels with target initial concentrations of 10^3^, 10^4^, or 10^5^ cells/ml and no OT1 cells. The surviving B16 cells were measured at 0, 24, 48, and 72 hours after inoculation into gels. The total number of data points *n* = 70 (**Supplemental Figure S1A**).
2. **Dataset 2** (short-term growth and killing): SIINFEKL-pulsed B16 melanoma cells were inoculated with target initial concentrations 10^4^, 10^5^, or 10^6^ cells/ml each with activated CD8^+^ OT1 cells with concentrations 0, 10^4^, 10^5^, 10^6^, or 10^7^ cells/ml. The surviving B16 cell numbers were measured at 0 and 24 hours. The total number of data points *n* = 175 (**Supplemental Figure S1B**).
3. **Dataset 3** (long-term growth and killing): SIINFEKL-pulsed B16 melanoma cells were inoculated with target initial concentrations 10^6^ or 10^8^ cells/ml each with OT1 T cells with concentrations 0, 10^6^, or 10^7^ cells/ml. Gels with B16 cell concentration of 10^8^ cells/ml were unstable, and thus was not included in the analysis. Measurements of surviving B16 cells were done at at 0, 24, 48, 72, and 96 hours post inoculation into gels. The total number of data points *n* = 96 (**Supplemental Figure S1C**).
4. **Dataset 4** (growth and killing in the first 24 hours): In this previously unpublished dataset, SIINFEKL-pulsed B16 melanoma cells were co-inoculated into gels with the target initial concentration of 10^5^ cell/ml and with OT1 T cells at the concentrations 0, 10^6^, or 10^7^ cell/ml. Surviving B16 cells were measured at 0, 4, 8, 12, and 24 hours post-inoculation into gels. The total number of data points *n* = 90 (**Supplemental Figure S1D**).
5. **Dataset 5** (killing at a high CTL concentration): In this previously unpublished dataset, SIINFEKL-pulsed B16 melanoma cells were co-inoculated into gels at the target initial concentration 10^5^ cells/ml and with OT1 cells at concentrations 0 or 10^8^ cells/ml. Surviving B16 cells were measured at 0 and 24 hours. The total number of data points *n* = 7 (**Supplemental Figure S1E**).

Experiments generating data for Datasets 1-4 were repeated three times (Experiments 1, 2, and 3), and each measurement was performed in duplicate [1]. These experimental duplicates were prepared for each experimental condition and at the specific time point each of the two gels was lysed, diluted, and the cells from each gel were plated into two 65 × 15 *mm*^2^ plates. Experiment generating Dataset 5 was performed once.

### Mathematical models

#### Mathematical models to explain tumor dynamics

Given previous observations of Budhu *et al.* [1] we assume that B16 melanoma (tumor) cells grow exponentially and are killed by OT1 CD8^+^ T cells (CTLs) at a rate proportional to the density of tumors. The change in the B16 cell concentration (*T*) over time is then described by a differential equation of the general form

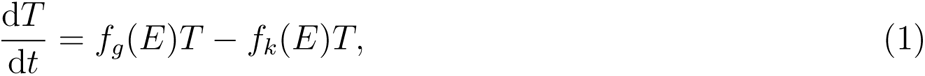

where *f_g_*(*E*) is the per capita growth rate and *f_k_*(*E*) is the death rate of tumors, and *E* is the CTL concentration. When *E* is constant, the general solution of this equation can be written as

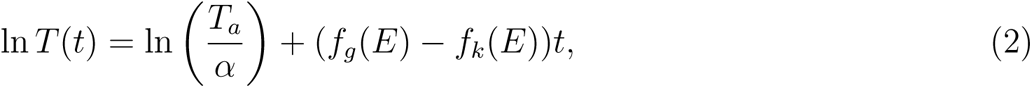

where we let *T* (0) = *T_a_/α* be the initial count which depends on the target B16 tumor cell concen- tration *T_a_* subject to a scaling factor *α*. It was typical to recover somewhat lower B16 cell numbers from the gel than it was targeted. For example, when targeting 10^4^ B16 tumor cells per ml in a gel it was typical to recover *∼* 4 × 10^3^ cells/ml at time 0 (e.g., **Supplemental Figure S1A**).

The simplest exponential growth and mass-action killing (**MA**) model assumes that tumors grow exponentially and are killed by CTLs at the rate proportional to CTL density (*f_g_*(*E*) = *r* and *f_k_*(*E*) = *kE*, **Figure 1A**). Using eqn. (2) change in the density of targets over time is given by

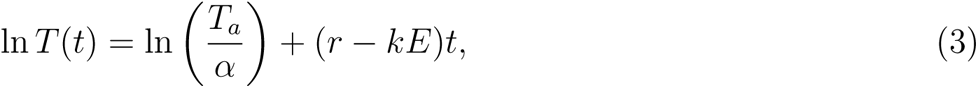

**Figure 1:**
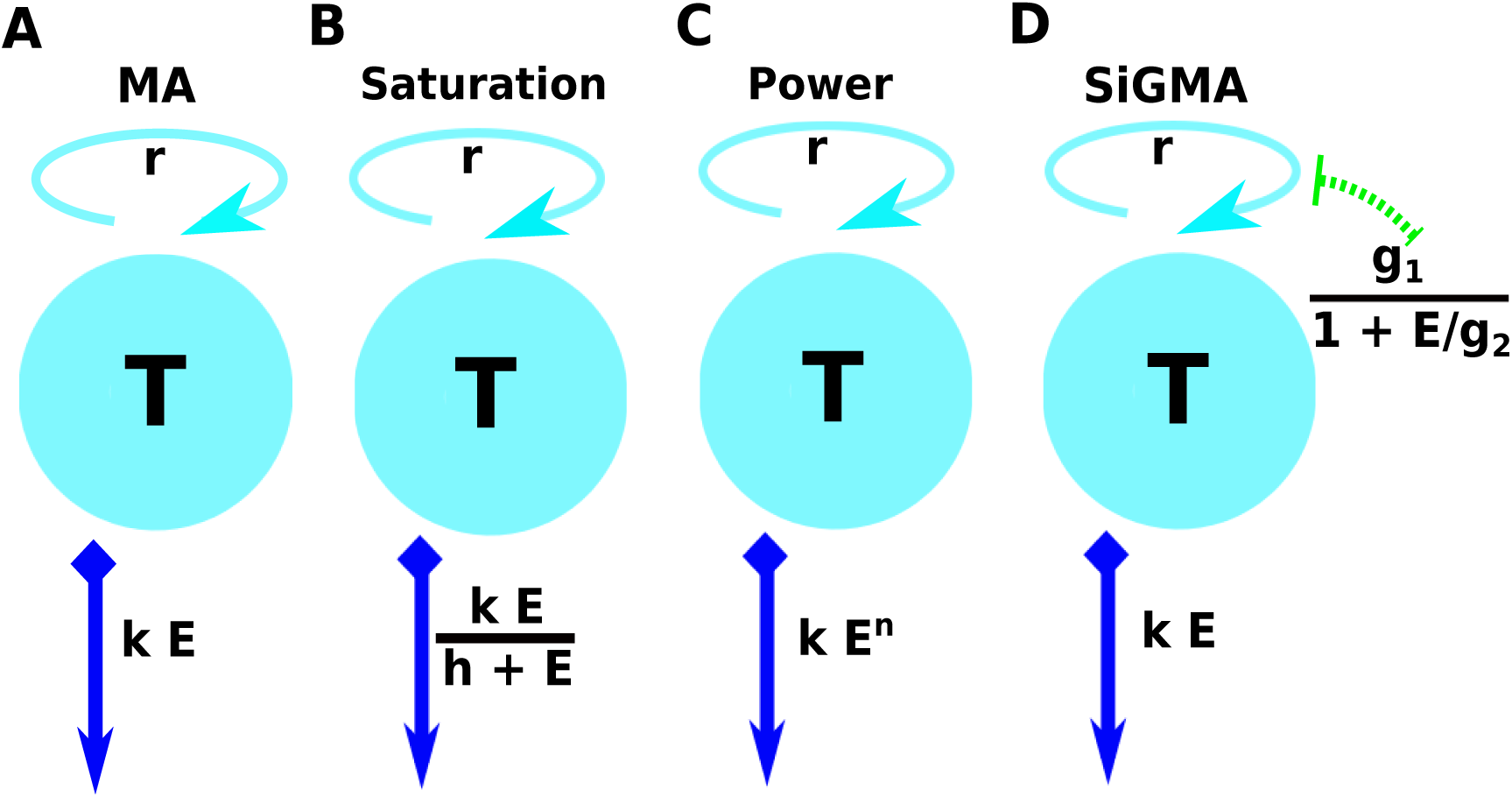
A schematic representation of the four main alternative models fitted to data on the dynamics of B16 tumor cells. These models are as follows: (A): an exponential growth of tumors and a mass-action killing by CTLs (**MA**) model (eqn. (3)); (B): an exponential growth of tumors and saturation in killing by CTLs (**Saturation** or **Sat**) model (eqn. (4)); (C) an exponential growth of tumors and killing by CTLs in accord with a powerlaw (**Power**) model (eqn. (5)); and (D) an exponential growth of tumors with CTL-dependent suppression of the growth and mass-action killing of tumors by CTLs (**SiGMA**) model (eqn. (6)). The tumor growth rate *r* is shown on the top of the cyan spheres which represent the B16 tumor cells *T* . For the suppression in growth model with a mass-action term in killing (D,“SiGMA”), the *E* dependent suppression rate is presented over the green arrow. The killing rate *k* for each model is shown in the blue arrow pointing downwards. For example, the Power model is shown by a constant growth rate *r* with the death rate of the tumors by *E* CTLs is *kE^n^*.

This model has three parameters (*r*, *k*, and *α*) to be estimated from the data.

The second “saturation” (**Sat**) model assumes that tumors grow exponentially and are killed by CTLs at a rate that saturates at high CTL densities 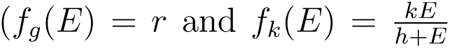, **Figure 1B**). Using eqn. (2) its solution is

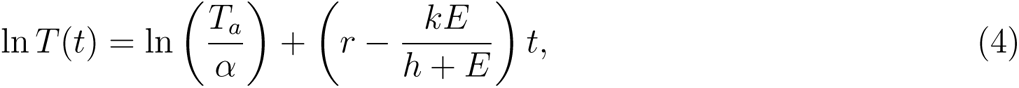

This model has 4 parameters (*r*, *k*, *h*, and *α*) to be estimated from the data.

The third “**Power**” model assumes that grow exponentially and are killed by CTLs at a rate that scales as a power law with CTL density (*f_g_*(*E*) = *r* and *f_k_*(*E*) = *kE^n^*, **Figure 1C**). Using eqn. (2) its solution is

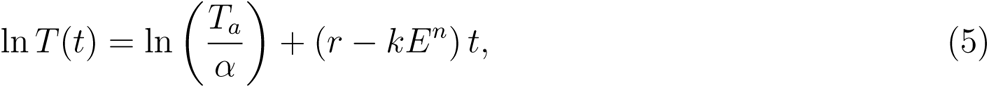

This model has 4 parameters (*r*, *k*, *n*, and *α*) to be estimated from the data.

In the fourth suppression-in-growth with mass-action-killing (**SiGMA**) model we assume that CTLs suppress growth rate of the tumor and kill the tumors according to mass-action law (*f_g_*(*E*) = 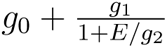 and *f_k_*(*E*) = *kE*, **Figure 1D**). Using eqn. (2) its solution is

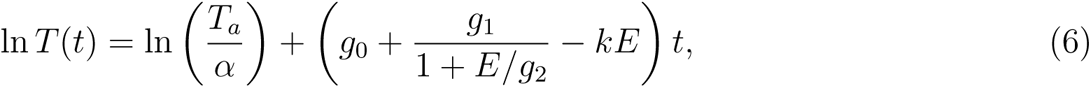

where *g*_0_ is the B16 tumor growth rate that is independent of the CTLs, and *g*_1_ the tumor growth rate that can be reduced by CTLs via non-lytic means, and *g*_2_ is the density of CTLs at which the growth rate *g*_2_ is reduced to half of its maximal value due to CTL activity. Note that in this model the rate of tumor cell replication in the absence of CTLs is *r* = *g*_0_ + *g*_1_. This model has 5 parameters (*g*_0_, *g*_1_, *g*_2_*, k*, and *α*) to be estimated from the data.

#### Estimating initial density of tumor cells in gels

In the general solution (eqn. (2)) we assumed that initial tumor density is proportional to the density targeted in experiments scaled by a factor *α*. We found that recovered concentrations of B16 tumor cells from gels at time *t* = 0 were consistently lower than the targeted value, and such reduction was approximately similar for different initial B16 concentrations (results not shown). Experimentally, this may arise because the clonogenic assay used to count the number of B16 tumor cells in the gels are not 100% efficient (results not shown). To check if assuming identical scaling factor *α* for the initial B16 concentration in different experiments we tested an alternative model where we assumed different *α* for different B16 cell concentrations. In this varying *α* model the first term in eqn. (2) can be written as ln 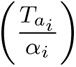 where *i* denotes the targeted B16 cell concentration. For example, a dataset containing targeted B16 concentrations of 10^5^, 10^6^ and 10^7^ cell/ml would have three *α*: *α*_1_, *α*_2_ and *α*_3_, respectively. For fitting the model with dataset-dependent *α* we used the function `MultiNonlinearModelFit` in Mathematica. We found that allowing *α* to vary between different targeted B16 concentrations when fitting the SiGMA model to Datasets 1-4 marginally improved the model fit 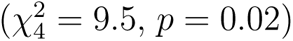 but did not influence estimates of other parameters (**Table S2**); in our following analyses we therefore opted for the simpler model with a single scaling parameter *α*.

#### Time to kill 90% of targets

To evaluate efficacy of CTL-mediated control of tumors we calculated the time it takes to kill 90% of tumors initially present. For every model (eqns. (3)–(6)) we solve an equation 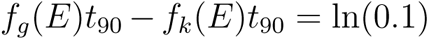 to find time *t*_90_ in terms of CTL concentration *E*:

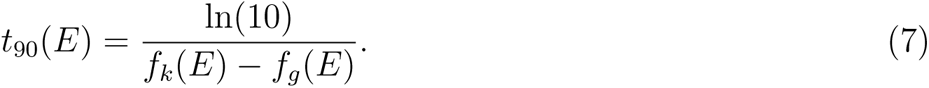

#### Models to explain tumor growth in the first 24 hours after inoculation into gels

In new experiments (Dataset 4) we found that growth of the tumors in the first 24 hours after inoculation into the gels may not follow a simple exponential curve. Experimentally, this delay may be due to the tumor cells adjusting to the gel environment. In order to explain this dynamics we propose two additional models. As a first alternative (**Alt 1**) model, we allow for a natural death of B16 tumor cells and then after a delay growth starts. The motivation for this new growth function comes from an algebraic sigmoid function which changes sign from a constant negative value to a constant positive value. The change in the concentration of B16 tumor cells in the absence of CTLs is given by

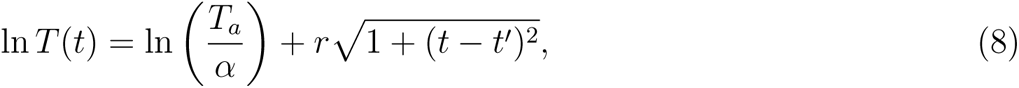

where the constant *t^′^* quantifies the time at which this change in sign happens. This model has 3 parameters (*α*, *r*, and *t^i^*) to be estimated from the data.

As the second alternative (**Alt 2**) model, we consider a mechanistic explanation of the non-linear dynamics of the tumor cells. We assume that a fraction *f_d_* of B16 tumor cells die at rate *d* and the rest (1 *− f_d_*) grow at rate *r*. The model can be described by the following equations

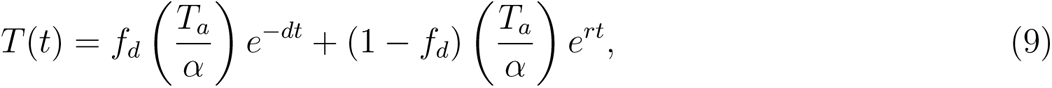

where *d* is the death rate of the *f_d_* subset of tumor cells. This model has 4 parameters (*α*, *f_d_*, *d*, and *r*) to be estimated from the data.

### Statistics

Natural log-transformed solutions of the models were fitted to the natural log of measured concentra- tions of B16 tumor cells using least squares. In the data there were 13 gels (out of 451) that had 0 B16 tumor cells recovered; these data were excluded from most of the analyses. The regression analyses were performed using function `NonlinearModelFit` Mathematica (ver 11.3.0.0). For every model we calculated AIC and ΔAIC as

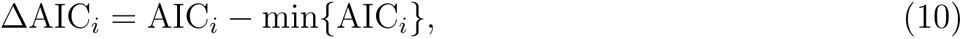

where subscript *i* denotes the model and ’min’ denotes the minimum for all the models [63]. The Akaike weight for the model *i* was calculated as

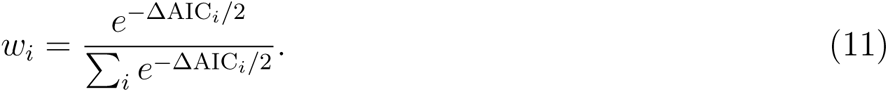

To evaluate the appropriateness of assumptions of least squares-based regressions we analyzed residuals of best fits by visual inspection and using Shapiro-Wilk normality test using the function ShapiroWilkTest in Mathematica.

## Results

### Experiments to measure how CTLs kill targets in collagen-fibrin gels

To estimate the den- sity of tumor-specific CTLs needed to control growth of B16 tumor cells, Budhu *et al.* [1] performed a series of experiments in which variable numbers of SIINFEKL peptide-pulsed B16 tumor cells and SIINFEKL-specific CTLs (activated OT1 CD8 T cells) were co-inoculated in collagen-fibrin gels and the number of surviving tumor cells was calculated at different time points (see **Supplemental Figure S1** and Materials and methods for more detail). In the absence of CTLs (**Dataset 1**), B16 tumor cells grew exponentially with the growth rate being approximately independent of the initial tumor density (**Supplemental Figure S1A**). Short-term (24 hours) experiments (**Dataset 2**) showed that when the density of CTLs exceeds 10^6^ cell/ml, the density of B16 cells declines in 24 hours, suggesting that the killing rate of the tumors exceeds their replication rate (**Supplemental Figure S1B**). Longer (96 hours) experiments (**Dataset 3**) showed that at high CTL densities (*>* 10^6^ cell/ml) the number of B16 targets recovered from gels declines approximately exponentially with time; interestingly, however, at an intermediate density of CTLs and B16 tumor cells of 10^6^ cells/ml, B16 cells initially decline but then rebound and accumulate (**Supplemental Figure S1C**). Previously unpublished experiments (**Datasets 4-5**) showed a similar impact of increasing CTL density on the B16 tumor dynamics during short-term (24h) experiments (**Supplemental Figure S1D–E**). Budhu *et al.* [1] concluded that the data from short- and long-term experiments (**Supplemental Figure S1A–C**) are consistent with the model in which the number of B16 tumor cells grows exponentially due to cell division and are killed by CTLs at a mass-action rate (proportional to the density of targets and CTLs). Budhu *et al.* [1] also concluded that density of 3.5 × 10^5^ cells/ml was critical for overall clearance of B16 tumor cells in collagen-fibrin gels.

### A simple exponential-growth-and-mass-action-killing model is not consistent with the data

The conclusion that a simple model with exponentially growing tumors and killing of the tumors by CTL via mass-action law (**MA** model, **Figure 1A**) was based on simple regression analyses of individual datasets (e.g., Dataset 1 or 3). To more rigorously investigate we proposed three additional models that made different assumptions of how CTLs impact B16 tumor cells including i) saturation in killing rate (**Sat** model, eqn. (4) and **Figure 1B**), ii) nonlinear change in the death rate of tumors with increasing CTL concentrations (**Power** model, eqn. (5) and **Figure 1C**), and iii) reduction in the tumor growth rate with increasing CTL concentrations and mass-action killing term (**SiGMA** model, eqn. (6) and **Figure 1D**). We then fitted these models including the MA model to all the data that in total includes 438 measurements (and excluded 13 gels with zero B16 tumor cells, see Materials and Methods for more detail). These data included two new unpublished datasets (Dataset 4 and 5) including the B16 tumor dynamics at physiologically high CTL concentrations (*E* = 10^8^ cell/ml, **Supplemental Figure S1E**). Interestingly, we found that the MA model fit these data with least accuracy while the Sat model (with a saturated killing rate) fitted the data best (**Supplemental Table S1**). Saturation in the killing rate by CTLs is perhaps not surprising in the full dataset given that in the Dataset 5 two gels inoculated with 10^5^ B16 tumor cells and 10^8^ cell/ml CTLs still contained B16 tumor cells at 24 hours (**Supplemental Figure S1E**). Because 10^8^ cells/ml is physiologically unrealistic density of CTLs in vivo, for most of our following analyses we excluded the Dataset 5.

**Table 1:** Parameters of the 4 alternative models fitted to Datasets 1-4 (excluding data with CTL = 10^8^ cell/ml) and metrics of quality of the model fits. Estimated parameters: α is a di- mensionless scaling factor, *r* is given in the units of per day, h is in cells/ml, g_1_ is in per day and g_2_ is in cells/ml. The parameter k have different units in different models: per OT1 cells/ml per day, per day, per OT1 (cells/ml)^n^ per day and per OT1 cells/ml per day for MA (eqn. (3)), Sat (eqn. (4)), Power (eqn. (5)) and SiGMA (eqn. (6)), respectively, and n is a dimensionless parameter. Parameter estimates and 95% confidence intervals for the best fit SiGMA model are: α = 2.71 (2.5 − 2.9), g_0_ = 0.12 (0.036 − 0.2)/day, g_1_ = 0.64 (0.55−0.73)/day, g_2_ = 6.72 (4.14−17.57)×10^3^ cell/ml, and k = 3.3 (3.15−3.4)×10^−7^ ml/cell/day; model fits are shown in Figure 2. The best fit model (with the highest w) is highlighted in blue.

| Datasets 1-4 ($E \leq 10^7$ cell/ml): n=431 | | | | | | | | | | | | |
| --- | --- | --- | --- | --- | --- | --- | --- | --- | --- | --- | --- | --- |
| Model | $\alpha$ | $r$, 1/day | $k$ | $h$ | $n$ | $g_0$ | $g_1$ | $g_2$ | SSR | AIC | ΔAIC | $w$ |
| MA | 2.77 | 0.576 | $3.79 \times 10^{-7}$ | | | | | | 164 | 814 | 160 | 0 |
| Sat | 2.81 | 0.744 | 6 | $5.34 \times 10^6$ | | | | | 118.5 | 677 | 23 | 0 |
| Power | 2.79 | 0.744 | $2.23 \times 10^{-4}$ | | 0.606 | | | | 116 | 668 | 14 | 0 |
| SiGMA | 2.71 | | $3.29 \times 10^{-7}$ | | | 0.12 | 0.64 | 6715 | 112 | 654 | 0 | 1 |

Importantly, the MA model was still the least accurate at describing the data from Datasets 1-4 which is visually clear from the model fits of the data as well as from statistical comparison of alternative models using AIC (**Figure 2** and **Table 1**). In contrast, the SiGMA model provided the best fit (**Table 1**). The SiGMA model is unique because it suggests that in these experiments CTLs impact tumor accumulation not only by killing the tumors but also by slowing down tumor rate of growth from the maximal value of *r* = *g*_0_ + *g*_1_ = 0.76/day to the minimal *g*_0_ = 0.12/day already at moderate CTL concentrations (*E ≈* 10^4^ cell/ml, **Table 1**). It is well recognized that CTLs are able to produce large amounts of interferon-gamma (**IFNg**) that may directly inhibit tumor growth, especially of IFNg-receptor expressing cells [64–66]. Interestingly, while statistically the Sat and Power models fit the data worse than the SiGMA model, visually the fits of these three models are very similar (**Figure 2**). Furthermore, at high CTL concentrations (*E* = 10^7^ cell/ml) all four models provide fits of a similar quality (**Figure 2E**).

**Figure 2:**
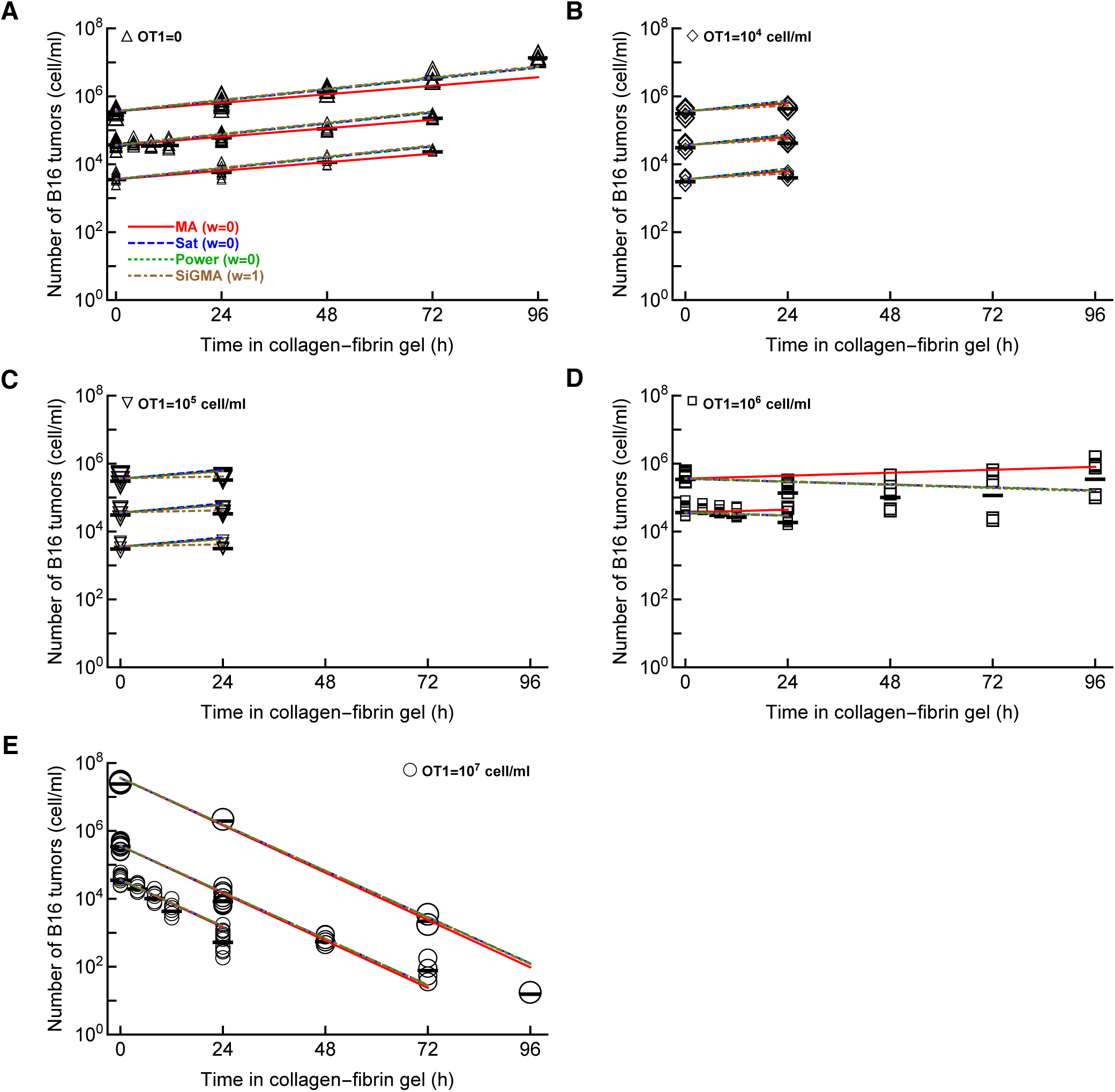
The model assuming exponential growth of B16 tumor cells and mass-action killing by CTLs is not consistent with the B16 tumor dynamics. We fitted mass-action killing (MA, eqn. (3) and Figure 1A), saturated killing (Sat, eqn. (4) and Figure 1B), powerlaw killing (Power, eqn. (5) and Figure 1C), and saturation in growth and mass-action killing (SiGMA, eqn. (6) and Figure 1D) models to data (Datasets 1-4) that includes all our available data with CTL densities *≤* 10^7^ cells/ml (see Materials and Methods for more detail). The data are shown by markers and lines are predictions of the models. We show model fits for data for (A): OT1 = 0, (B): OT1 = 10^4^ cell/ml, (C): OT1 = 10^5^ cell/ml, (D): OT1 = 10^6^ cell/ml, and (E): OT1 = 10^7^ cell/ml. Parameters of the best fit models and measures of relative model fit quality are given in Table 1; Akaike weights *w* for the model fits are shown in panel A.

It is important to note that even the best fit SiGMA model did not accurately describe all the data. For example, the model over-predicts the B16 counts at 24 hours for OT1 concentrations 10^4^ and 10^5^ cells/ml (**Figure 2B–C**) and under-predicts the B16 counts at 96 hours in growth (**Figure 2A**) and at 72 hours for OT1 concentrations 10^7^ cells/ml (**Figure 2E**).

To intuitively understand why the MA model did not fit the data well we performed several regres- sion analyses. Specifically, for every CTL and B16 tumor cell concentrations we calculated the net growth rate of the tumors *r*_net_ (**Figure S1**); in cases of several different targeted B16 concentrations we calculated the average net growth rate. In the absence of CTLs, the net growth rate of tumor cells was *r*_0_ = 0.62/day (**Figure S2**). Then for every CTL concentration we calculated the death rate of B16 tumor cells due to CTL killing as *K* = *r*_0_ *− r*_net_. For the MA model, the death rate *K* should scale linearly with the CTL concentration [42], however, we found that this was not the case for B16 tumor cells in gels where the death rate scaled sublinearly with the CTL concentration (**Figure S2**). Importantly, this analysis also illustrates that at low CTL concentrations (10^4^ *−* 10^5^ cell/ml) we observe a much higher death rate of targets than expected at the power *n* = 0.57 (**Figure S2**). This indirectly supports the SiGMA model that predicts a higher (apparent) death rate of targets at low CTL concentrations due to reduced tumor’s growth rate.

One feature of these experimental data is that the recovery of the B16 tumor cells from the gels was typically lower than the targeted concentration that somewhat varied between different experiment and targeted B16 cell numbers (e.g., **Supplemental Figure S1**). Instead of fitting individual parameters to estimate the initial density of B16 tumor cells for every target B16 concentration we opted for an alternative approach. To predict initial concentration of B16 tumor cells we fitted a parameter *α* that scaled the targeted B16 number to the initial measured B16 concentration in the gel (see Materials and Methods for more detail). In separate analyses we investigated if assuming different *α* for different target B16 tumor concentrations by fitting our best fit models (for Datasets 1-4 or Datasets 1-5) with one or 5 *α* (see Materials and methods for more detail and **Supplemental Table S2**). Interestingly, the SiGMA model with varying *α* fitted the data (Datasets 1-4) marginally better than model with one *α* (F-test for nested models, *p* = 0.02, **Supplemental Table S2**). Other parameters such as the B16 tumor growth rate and CTL kill/suppression rates, however, were similar in both fits (**Supplemental Table S2**). In contrast, the fits of the Sat model to all data (Datasets 1-5) were similar whether we assumed different or the same *α* for different target B16 tumor concentrations (*p* = 0.37, **Table S2**). Because in all cases other statistical features of the model fit (e.g., residuals) were similar, in most of the following analyses we considered a single parameter *α* in fitting models to the data.

In our datasets we had in total 13 gels which did not contain any B16 tumor cells after co-incubation with CTLs (**Supplemental Figure S1B&C**); these data were excluded from the analyses so far. Data exclusion may generated biases, and we therefore investigated if instead of 0 B16 targets we assume these measurements are at the limit of detection (**LOD**). The true limit of detection was not defined in these experiments so we ran analyses assuming that LOD = 2 *−* 10 cell/ml. Importantly, inclusion of these 13 gels at the LOD did not alter our main conclusion; specifically, the SiGMA model remained the best model for Datasets 1-4 and the Sat model remains the best model when we used Datasets 1-5 (results not shown).

### The best fit model varies with chosen subset of the data

Experimental data sug- gest that the CTL (OT1) concentration of approximately 10^6^ cell/ml is critical for removal of B16 melanoma cells [1]. Specifically, at concentrations *E <* 10^6^ cell/ml the tumor cell concentration increases (**Supplemental Figure S1A–C**) while at *E >* 10^6^ cell/ml tumor cell concentration declines (**Supplemental Figure S1B–D**). The growth and death rates of the tumors are similar when *E ≈* 10^6^ cells/ml and interestingly, in one dataset, the B16 tumor concentration initially declines but after 48h starts to increase (**Supplemental Figure S1C**). None of our current models could explain this latter pattern. To investigate if the data with CTL concentrations of 10^6^ cell/ml may bias the selection of the best fit model we fitted our 4 alternative models to the data that excludes gels with B16 target concentrations of 10^5^ and 10^6^ cell/ml and CTL concentrations of 10^6^ cell/ml from Dataset 3 and Dataset 4, respectively. Interestingly, for these subset of data the Power model fitted the data with best quality (based on AIC) predicting that the death rate of B16 tumor cells scales sub-linearly (*n* = 0.42) with CTL concentration (**Supplemental Table S3**). The Power model also provided the best fit if we included 7 additional gels from the Dataset 5 (with highest CTL concentrations, **Supplemental Table S3**). Interestingly, the MA model fitted this data subset with much better quality visually even though statistically the fit was still the worst out of all four models tested (**Supplemental Table S3** and results not shown).

We further investigated if focusing on smaller subsets of data may also result in other models fitting such data best. For example, in one approach we focused on fitting the models to subsets of data with a single target B16 tumor cell concentration (**Supplemental Table S4**). Interestingly, for B16 concentrations of 10^4^ and 10^6^ cell/ml, the Power model provided the best fit but for target B16 concentration of 10^5^ cell/ml, the Power and SiGMA models gave best fits. Including Dataset 5 in these analyses often led to the Sat model being the best (**Supplemental Table S4**). Finally, dividing the data into subsets for different experiments (out of 3), the Power model fitted best the data from Experiment 1 and 3 and SiGMA model fitted best the data from Experiment 2 (see **Supplemental Table S5**). Taken together these analyses strongly suggest that selecting the best model describing the dynamics of B16 tumor cells depends on the specific subset of data chosen for the analysis.

### Alternative models predict different CTL concentrations needed to control tumor growth

Given the difficulty of accurately determining the exact model for B16 tumor growth and its control by CTLs one could wonder why we need to do that. To address this potential criticism we calculated the time (eqn. (7)) it would take for CTLs to eliminate most (90%) of tumor cells if CTLs control tumor growth in accord with one of the 4 alternative models (e.g., with parameters given in **Table 1**). Interestingly, the MA model predicted the largest CTL concentration that would be required to eliminate most of the tumor cells in 100 days while the SiGMA model required the fewest (1.54 × 10^6^cell/ml vs. 0.41 × 10^6^cell/ml, respectively, **Figure 3**). The 4-fold difference may be clinically substantial in cancer therapies using adoptively transferred T cells (e.g., in tumor infiltrating lymphocyte-based therapies [67]). Interestingly, however, that the difference in predicted CTL concentration was somewhat similar for SiGMA and Power models that provided best fits for subsets of the data (**Figure 3**). Interestingly, the range of CTL concentrations was wider between alternative models fitted to subsets of the data (results not shown) further highlighting the need of better, more rigorous understanding how CTLs control tumor’s growth in collagen-fibrin gels.

**Figure 3:**
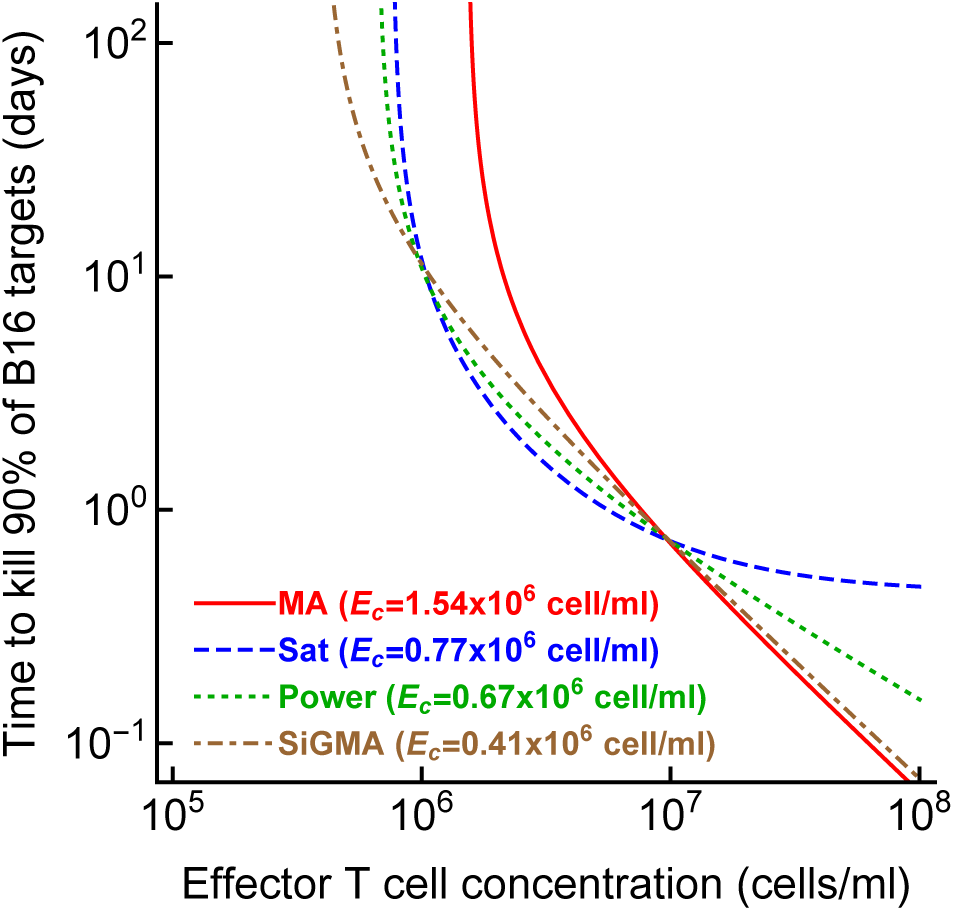
The CTL concentration needed to eliminate most B16 tumor cells depends on the model of tumor control by CTLs. For every best fit model (**Table 1**) we calculated the time to kill 90% of B16 targets for a given concentration of CTLs (eqn. (7)). For every model we also calculated the control CTL concentration (*E_c_*) that is required to eliminate at least 90% of the tumor cells within 100 days.

### Mathematical models different from simple exponential growth are needed to explain B16 tumor dynamics in the absence of CTLs

In our analyses so far we focused on different ways CTLs can control growth of the B16 tumor cells which assuming that in the absence of CTLs tumors growth exponentially (eqns. (3)–(6)). In our new Dataset 4 in which gels were sampled at 0, 4, 8, 12, and 24 hours after inoculation we noticed that B16 tumor cells did not grow exponentially early after inoculation into gels (**Supplemental Figure S1D**). We therefore investigated whether a simple model in which B16 tumor cells grow exponentially is in fact consistent with our data.

First, we fitted the exponential growth model (eqn. (3) with *E* = 0) to all data from Dataset 1-5. Interestingly, while the model appeared to fit the data well (**Figure 4A**) and statistically the fit was reasonable (e.g., residuals normally distributed), model fits did not describe all the data accurately. In particular, the model over-predicted the concentration of B16 tumor cells at low (10^3^ *−*10^4^ cell/ml) and high (10^8^ cell/ml) targeted B16 concentrations. Lack of fit test also indicated that the model did not fit the data well (*F*_20,154_ = 7.12, *p <* 0.001). Finally, allowing the tumor growth rate to vary with the targeted B16 concentration resulted in a significantly improved fit (*F*_4,170_ = 19.77, *p <* 0.001) suggesting that the growth rate of B16 tumor cells in the absence of CTLs may be density-dependent (*r*_0_ = 0.59/day, *r*_0_ = 0.65/day, *r*_0_ = 0.64/day, and *r*_0_ = 0.85/day, *r*_0_ = *−*0.15/day for targeted B16 tumor cell concentrations of 10^3^, 10^4^, 10^5^, 10^6^, and 10^7^ cell/ml, respectively, and *α* = 2.48).

**Figure 4:**
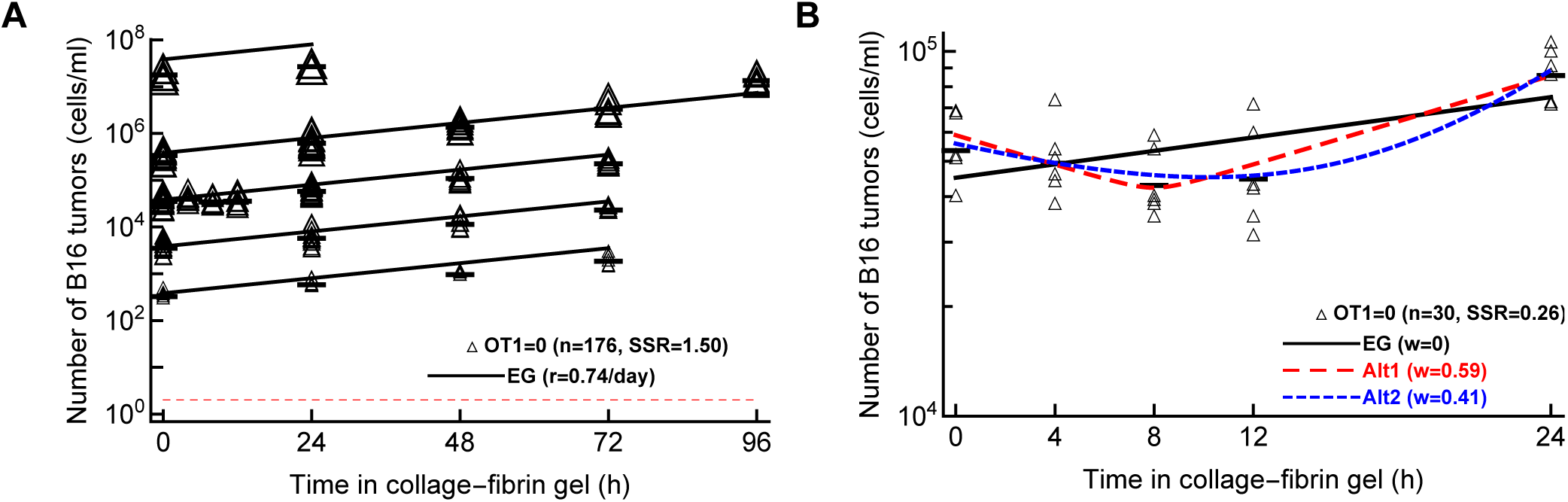
Pure exponential growth (EG) model is not consistent with the data on B16 tumor dynamics in the absence of CTLs. (A): we fitted with an exponential growth model (eqn. (3) with *E* = 0) to data on B16 growth from all datasets 1-5 with OT1 = 0. The best fit values for the parameters along with 95% confidence intervals are: *α* = 2.6 (2.4 *−* 2.8) and *r* = 0.74 (0.69 *−* 0.79)/day. (B): we fitted exponential growth and two alternative models (eqn. (3) with *E* = 0 and eqns. (8)–(9)) to the data from Dataset 4 for which OT1 = 0. The relative quality of the model fits is shown by Akaike weights *w* (see **Table S6** for model parameters and other fit quality metrics). The data are shown by markers and model predictions are shown by lines.

Second, we noticed that in our new dataset with B16 tumor growth kinetics recorded in the first 24 hours after inoculation into gels (Dataset 4) does not follow a simple exponential increase (**Supplemental Figure S1D**). Instead, there is appreciable decline and then increase in the B16 cell concentration. We therefore fitted an exponential growth (**EG**) model along with two alternative models that allow for non-monotonic dynamics – i) a phenomenological model (eqn. (8)), and ii) a mechanistic model allowing for 2 sub-populations of tumor cells, one dying and another growing over time (eqn. (9)). Interestingly, while the EG model did not fit the data well, either of the alternative models described the data relatively well (**Figure 4**). These analyses thus strongly suggest that the dynamics of B16 tumor cells in collagen-fibrin gels in the absence of CTLs are not consistent with a simple exponential growth model.

### Experiments with several measurements of B16 tumor concentrations at specifically chosen CTL densities will best allow to discriminate between alternative models

In several alternative analyses we found that the best model describing the dynamics B16 tumor cells in collagen fibrin gels depends on specific dataset chosen for the analysis. It is unclear why this may be the case. One potential explanation is that individual datasets are not balanced, some have more measurements but on a shorter time scale while others are of a longer duration with fewer replicates. Because the exact mechanism of how CTLs impact tumor dynamics is important in predicting the concentration of CTLs needed for tumor elimination (**Figure 3**), we next sought to determine whether specific experimental designs may be better suited to discriminate between alternative models [62]. We therefore performed stochastic simulations to generate “synthetic” data from a given assumed model for different experimental designs and tested whether by fitting alternative models to the synthetic data we can recover the model used to generate the data.

We considered three different designs and compared two types within each design.

- **Design D1**: Two time point experiment (Type A) vs four time point experiment (Type B). The two time point experiment have 48 observations. B16 target concentrations are 10^3^, 10^4^, 10^5^, 10^5^, 10^6^, 10^7^, 10^8^ cells/ml, OT1 concentrations are 0, 10^5^, 10^6^, 10^7^ cells/ml and time points are 0 and 24 hours. The four time point experiment have 48 observations. B16 target concentrations are 10^5^, 10^6^, 10^7^ cells/ml, OT1 concentrations are 0, 10^5^, 10^6^, 10^7^ cells/ml and time points are 0, 24, 48, 72 hours.
- **Design D2**: Short-term experiment (Type A) vs long-term experiment (Type B). The short time experiment have 48 observations. B16 target concentrations are 10^5^, 10^6^, 10^7^, OT1 concentrations are 0, 10^5^, 10^6^, 10^7^ cells/ml and time points are 0, 8, 16, 24 hours. The long time experiment have 48 observations. B16 target concentrations are 10^5^, 10^6^, 10^7^ cells/ml, OT1 concentrations are 0, 10^5^, 10^6^, 10^7^ cells/ml, and time points are 0, 24, 48, 72 hours.
- **Design D3**: More frequent OT1 experiment (Type A) vs less frequent OT1 experiment (Type B). The more frequent OT1 experiment have 40 observations. B16 target concentrations are 10^5^, 10^6^ cells/ml, OT1 concentrations are 0, 5 × 10^5^, 10^6^, 5 × 10^6^, 10^7^ cells/ml and time points are 0, 24, 48, 72 hours. The less frequent OT1 experiment have 40 observations. B16 target concentrations are 10^5^, 10^6^ cells/ml, OT1 concentrations are 0, 10^4^, 10^5^, 10^6^, 10^7^ cells/ml and time points are 0, 24, 48, 72 hours.

To draw a statistical comparison between the Types A and B of the experimental designs described above, we first chose one of the Saturation, Power, or SiGMA models with their best fit parameters (**Table 1**) and generated 48 observations for D1 and D2, or 40 observations for D3 for each of Types A and B. We excluded the MA model from these analyses as it never fitted the data well. In each of the generated predictions we added an error randomly chosen from the list 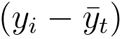, where *y_i_* is the observed B16 count in the data and 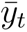 is the average of *y_i_* at time *t*. Next, we simulated 100 replicates of such pseudo experiments, fitted the three models (Sat, Power, and SiGMA) to these 100 replicates, and computed the Akaike weights to determine the best fit model for each replicate. Due to the randomly chosen error structure for these hypothetical experiments, we found substantial variability among these 100 replicates where the best fit model was often different from the model from which the identical replicates were generated. For example, generating 100 simulated datasets from the Saturation model, we found that the Saturation model fitted these data only in 52% cases while the Power model fitted the best 36% of the time and the SiGMA model 12% of the time (first column of the first Type A matrix of **Supplemental Figure S5D1**).

By repeating the analysis for all three models we generated a matrix of Akaike weights with diagonal terms being heavier than the off-diagonal terms along with a constraint that the sum of a column should always add up to one (see **Supplemental Figure S5**). In this representation, a better experimental design among each types has a heavier diagonal than off-diagonal elements. Following this rule we see that D1 (Type B), D2 (Type B) and D3 (Type A) are the better experiment types (**Supplemental Figure S5**). To show that the difference between experimental design are statistically significant we used a resampling approach. We defined a test statistic measure given by

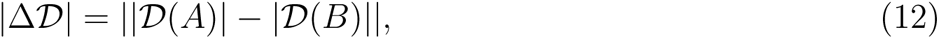

where *Δ* is the determinant of the matrix and |Δ*Δ|* is the absolute difference between two deter- minants. |Δ*Δ|* is equivalent to a difference in volume of two 3D parallelepipeds which edges are the columns of a matrix. For the hypothesis testing we defined the null hypothesis as: the column vectors, with constraints that the sum of elements must be unity, belong to the same class for both the experimental designs. We performed a null distribution test and a permutation test to reject the null hypothesis and showed that the column vectors which constitute the experimental designs are significantly different.

For the null distribution test, we randomly generated Type A and Type B sets of 10^6^ matrices with their columns being normalized to unity. |Δ*Δ|* was then computed for the the Types A and B which forms a universal null distribution. The p-value is then the number of times |Δ*Δ|*_null_’s are greater than observed |Δ*Δ|*_obs_ normalized by the total number of simulations (10^6^). The p-values for each of designs D1, D2 and D3 (**Supplemental Figure 5B**) confirms that a long time experiment with more time point observations and closely spaced CTL concentrations is a significantly better experimental design. For the permutation test, we generated three column matrices from all permutations of the six columns for each of designs D1, D2 and D3. The columns were chosen from the constructed matrices of **Supplemental Figure S5**. Then we randomly chose sets of two matrices for Types A and B from all the permutations of the previous step. |Δ*Δ|*_per_ was computed for the Types A and B which forms a distribution. The p-value was then the number of times the permuted |Δ*Δ|*_per_’s are greater than observed |Δ*Δ|*_obs_ normalized by the total number of permuted sets (**Supplemental Figure S5**). With a permutation test we found that a long time experiment with more time point observations is a significantly better experiment but fail to confirm the same for closely spaced OT1 concentrations with statistical significance (see right panels of **Supplemental Figure S5** for p-values). Taken together, these simulations suggest that longer experiments with at least 4 time points and a variable CTL concentration should provide best statistical power to discriminate between alternative models of B16 tumor control.

## Discussion

Quantitative details of how CTLs kill their targets in vivo remain poorly understood. Here we analyzed unique data on the dynamics of SIINFEKL peptide-pulsed B16 melanoma tumor cells in collagen-fibrin gels – that may better represent in vivo tissue environments — in the presence of known numbers of SIINFEKL-specific CTLs (OT1 T cells) [1]. We found that a previously proposed model in which tumors grow exponentially and are killed by CTLs proportional to the density of CTLs (mass-action law) did not describe the experimental data well. In contrast, the model in which CTLs suppress the rate of tumor replication and kill the tumors in accord with mass-action law fitted a subset of the data (Datasets 1-4 with physiologically relevant CTL concentrations of *E ≤* 10^7^ cell/ml with best quality (**Table 1**). This result raises an interesting hypothesis that control of tumors by CTLs may extend beyond direct cytotoxicity, e.g., by secretion of cytokines. In fact, previous observations suggested that IFNg and TNFa may suppress tumor growth in different conditions although the ultimate effect of these cytokines on tumor progression in vivo is inconclusive as IFNg may in fact improve metastasis of some tumors [64–66].

Importantly, however, fitting the alternative models to different subsets of data resulted in dif- ferent best fit models, e.g., including the data with high CTL concentrations (*E ≤* 10^8^ cell/ml) typically predicted that the death rate of B16 tumor cells saturated at high CTL concentrations (**Supplemental Table S1**). In other cases, a power model in which the death rate of tumors scales sublinearly with the CTL concentration described subsets of the data best (**Supplemental Table S3**). Analysis of a new dataset on B16 tumor growth in the first 24 hours after inoculation into gels with no CTLs suggested that a simple exponential model does not describe these data adequately; instead models that allow for initial loss and then rebound in the number of B16 tumor cells was the best (**Figure 4B**). We also developed a novel methodology and proposed designs of experiments that may allow to better discriminate between alternative mathematical models. Our analysis sug- gested that longer-term experiments (0-72 hours) with 4 measurements of B16 cell concentration with several OT1 concentrations would have the highest statistical power (**Figure 5**).

**Figure 5:**
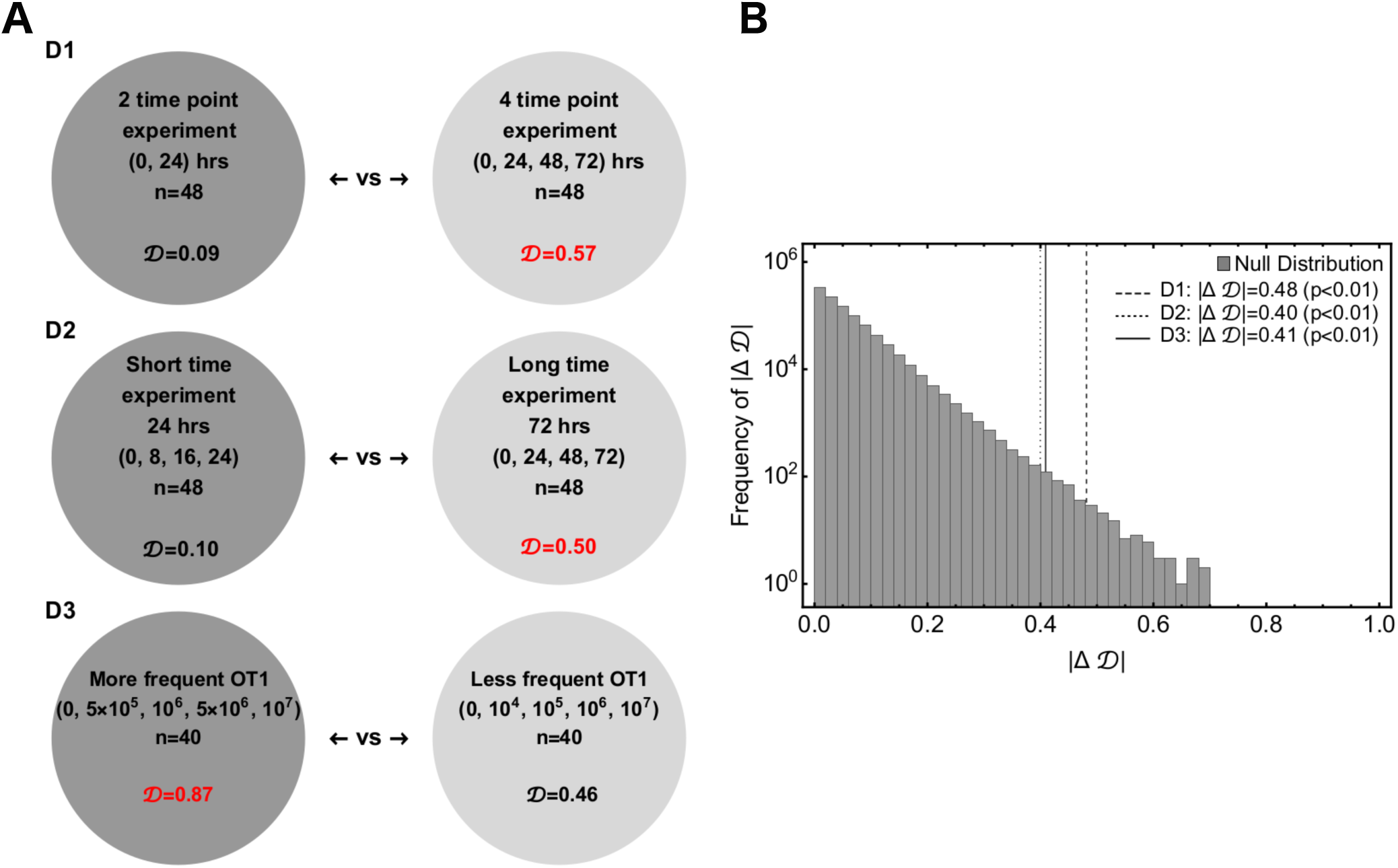
Power analysis indicates that longer experiments with several, closely spaced CTL concentrations would allow to best discriminate between alternative models. We performed three sets of simulations to get insights into a hypothetical future experiment which may allow to better discriminate between alternative mathematical models. (A): Three experimental designs are: D1 – 2 time point vs 4 time point experiments; D2 — short time scale (0-24h) vs. long time scale (0-72h) experiments; D3 — more frequently chosen values of CTL concentrations vs less frequently chosen vales of CTL concentrations (see Figure S5 and Materials and methods for more details). For every experimental setup we calculate *Δ* – the determinant of a matrix formed from a simulated experimental set whose columns are constrained. (B): We define a test measure |Δ*Δ|*_obs_ between two sets of each of D1, D2 and D3 and compare the observed |Δ*Δ|*_obs_ with the universal null distribution of |Δ*Δ|*_null_ to compute the p-value. The values of *Δ* in red in panel A shows the better experimental designs in the pairs.

Determining the exact mechanism by which CTLs control growth of B16 tumors may go beyond academic interest. In T cell-based therapies for the treatment of cancer, knowing the number of T cells required for tumor control and elimination is important. Our analysis suggests that specific details of the killing term do impact the minimal CTL concentration needed to reduce the tumor size within a defined time period (**Figure 3**). Other parameters characterizing impact of CTLs on tumor growth may also be important (**Figure 6**). For example, our analysis suggests that tumor’s growth rate, per capita killing rate by CTLs or the overall death rate of the tumors depend differently on CTL concentration given the underlying model (**Figure 6A–C**). The latter parameter, the death rate of CTL targets, has been estimated in several previous studies (reviewed in [44]) and ranges from 0.02/day to 500/day [37, 39, 40, 43, 68–72]. While our estimates are consistent with this extremely broad range whether killing of B16 tumor cells in collagen-fibrin gels occurs similarly to elimination of targets in vivo (peptide-pulsed or virus-infected cells) remains to be determined. Interestingly, our models predict a highly variable number of B16 tumor cells killed per day especially at low CTL concentrations (**Figure 6D**). We estimate that a relatively small number of targets are killed per CTL per day that is in line of previous estimates for in vivo killing of peptide-pulsed targets by effector or memory CD8 T cells [42, **Figure 6D**].

**Figure 6:**
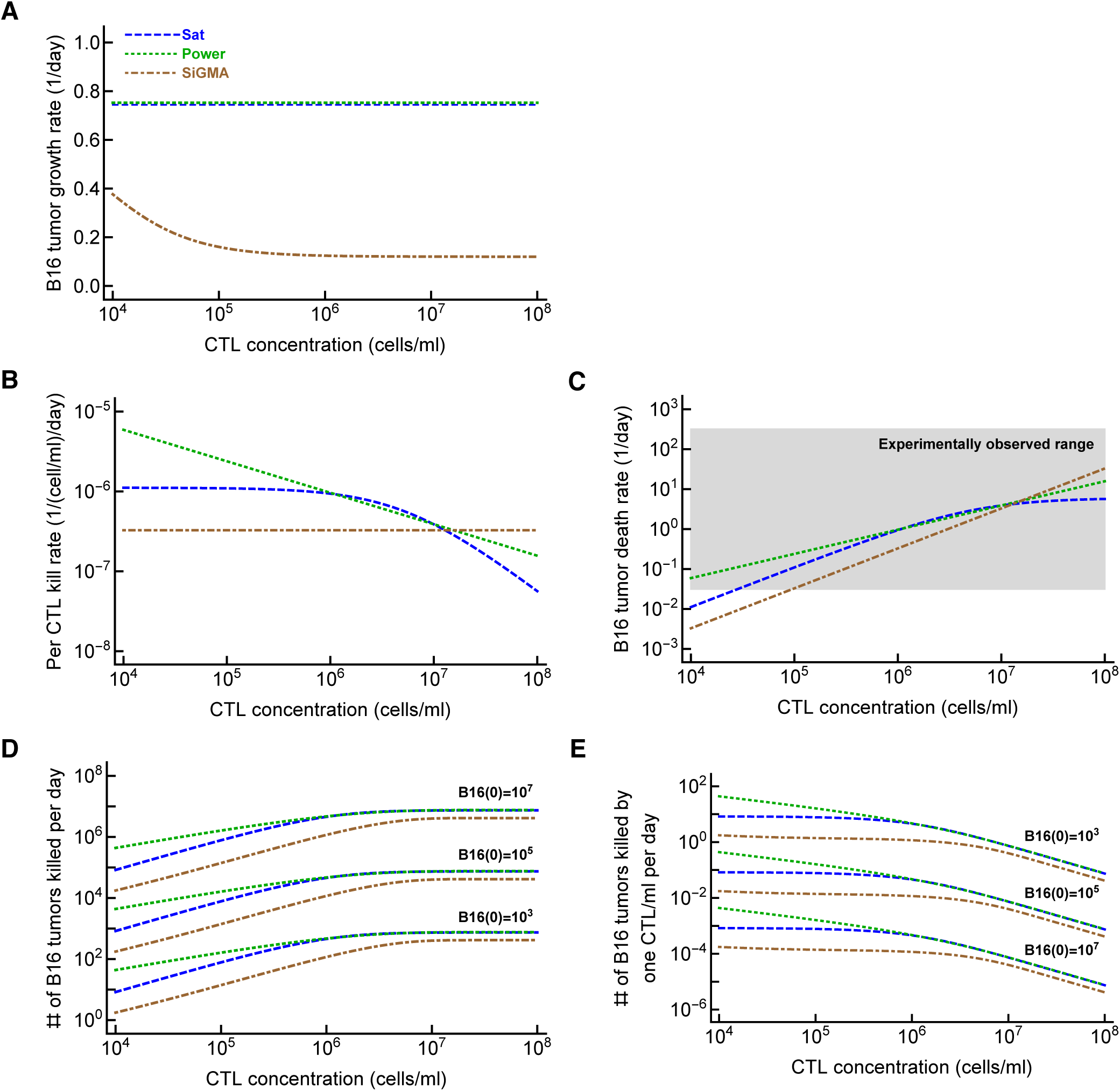
Metrics to quantify efficacy of CTL-mediated control of tumors are model-dependent. For the three alternative models (Sat, Power, and SiGMA) that fitted some subsets of data with best quality we calculated metrics that could be used to quantify impact of CTLs on tumor growth depending on the concentration of tumor-specific CTLs. These metrics include (A): the growth rate of the tumors (*f_g_* in eqn. (1)); (B): per capita kill rate of tumors (per 1 CTL per day, *f_k_/E* in eqn. (1)); (C): the death rate of tumors due to CTL killing (*f_k_* in eqn. (1)). The grey box shows the range of experimentally observed death rates of targets as observed in some previous experiments (see Discussion for more detail and [44]); (E): the total number of tumors killed per day as the function of 3 different initial tumor cell concentrations (indicated on the panel); and (D): the number of tumors killed per 1 CTL/ml per day. The latter two metrics were computed by taking the difference of growth and combined killing at 24 hours. The parameters for the models are given is **Table 1** and model equations are given in eqns. (4)–(6).

Our work has several limitations. First, specifics of tumor cell and CTL movements in the gels remain poorly defined. Previous studies suggested that CTL motility in collagen-fibrin gels may be anisotropic creating bias in how different CTLs locate their targets [61]. Second, errors in estimating the number of surviving B16 tumor cells have not been quantified. For example, in some cases zero B16 cells were isolated from the gels while other gels in the same conditions contained tens-to- hundreds of cells (**Supplemental Figure S1B–C**). In our experience, the clonogenic assays typically do not allow to recover 100% of inoculated cells that is also indicated by estimated parameters *α >* 1. In fact, *α* = 2.8 suggests that only 1/2.8 = 35% of inoculated B16 tumor cells are typically recovered.

In the way of how we fitted models to data (by log-transforming model predictions and the data), we had to exclude the gels with zero B16 tumor cells from the analysis. While this exclusion did not impact our overall conclusions, future studies may need to develop methods to include 0 values in the analysis. Third, the density of gels may change over the course of experiment reducing the ability of CTLs to find their targets. Using microscopy to track tumor cells and CTLs may better define if the movement patterns of the cells change over time in the gel. Fourth, the dynamics of CTLs and loss of peptide by B6 tumor cells have not been accurately measured. In particular, we observed that at CTL concentration of 10^6^ cells/ml and targeted B16 tumor cell concentration of 10^6^ cell/ml, after the initial decline, B16 tumor concentration rebounded (**Figure S1C**). Decline in CTL concentration with time could be one explanation; however, in other conditions, B16 tumor cells continue declining exponentially, arguing against a loss of CTLs in the gels. Tumor escape could be another explanation. Future experiments would benefit from also measuring CTL concentration in the gel, along with B6 tumor cells, especially in longer (48-72h) experiments. Fifth, the final fits of the models to data did not pass the assumption of normality as the residuals were typically not-normally distributed (e.g., by Shapiro-Wilk normality test). We have tried several methods to normalize the residuals (e.g., excluding the outliers, using arcsin(sqrt) transformation) but none worked. Whether non-normal residuals led to biased parameter estimates of our best fit models remains to be determined. Sixth and finally, we assumed that every CTL is capable of killing and every target is susceptible to CTL-mediated killing which may not be accurate. Indeed, the result that Power model fits several subsets of data with best quality and predicts sublinear increase in the death rate of targets with CTL concentration may be due to heterogeneity in CTL killing efficacy. However, such a model would need to assume that inoculation of CTLs into gels results in a bias of inoculating a smaller fraction of killer T cells at higher CTL concentrations which seems unlikely.

Our work opens up avenues for future research. One curious observation of Budhu *et al.* [1] is that the death rate of B16 tumor cells does not depend on the concentration of the targets in the gel. We confirmed this observation as the models that include dependence of the B16 tumor cell death rate on tumor cell concentration (e.g., the updated SiGMA model with *f_k_* = *kE/*(1 + *a*_1_*T* + *a*_2_*E*) did not improve the fit quality (e.g., in the best fits of Datasets 1-4 we found *a*_1_ *→* 0 and *a*_2_ *→* 0). This model-driven experimental observation is inconsistent with effector to target ratio-dependence in chromium release assays and with many theoretical arguments suggesting that killing of targets (or interactions between predators and preys) should be ratio-dependent, not density-dependent [29, 73–75]. Interestingly, our analysis of data from experiments on killing of peptide-pulsed targets in murine spleens by activated and memory CD8 T cells also showed no dependence on target cell concentration [42]. Future studies need to reconcile the difference between theoretical arguments and in vitro experiments and experimental observations in gels and in vivo.

The hypothesis that CTLs may impact the rate of tumor growth in collagen-fibrin gels can be tested experimentally. One such experiment could be to use two populations of tumors expressing different antigens, e.g., SIINFEKL and Pmel, in the presence or absence of SIINFEKL-specific CTLs (OT-1 T cells) [76]. Our experiments and mathematical modeling-based analyses can be extended to other types of tumor cells, CTL specifitiy, and the type of gels. Whether the CTL killing rates estimated from in vitro data correlates with CTL efficacy in vivo remains to be determined. Effective cancer immunotherapy relies on the infiltration and killing response of CD8^+^ T-cells [77, 78]. Increase of intratumoral CD8^+^ T-cells are shown to have direct correlation with radiographic reduction in tumor size in patients responding to treatment [79]. In B16 preclinical melanoma models cancer vaccines are found to induce cancer specific CD8^+^T-cells into tumors leading to cytotoxicity [80]. Estimating CTL killing efficiency such as kill rate per day or the number of melanoma cells killed per day could be useful in providing guidelines on cancer immunotherapy research and thus our modeling platform could therefore provide valuable insights for estimating the efficacy of T-cell based immunotherapies against cancer. The collagen-fibrin platform could be also useful to determine the killing efficiency of T cells (either expanded tumor infiltrating lymphocytes (TILs) or chimeric antigen receptor (CAR) T cells) prior to adoptively transferring them into patients; correlating this killing efficacy metric with actual success or failure of the therapy in patients may be a cheaper way to predict the overall efficacy of the therapy thus saving time and resources.

### Data sources

The data for the analyses is provided as a supplement to this publication and on github: https://github.com/vganusov/killing_in_gels.

## Abbreviations

CTLs: cytotoxic T lymphocytes
MA: mass action
Sat: saturation
SiGMA: suppression in growth with mass action in killing.

## Code sources

All analyses were performed in Mathematica. The sample code of fitting alternative models to the data is provided as a supplement to the paper and on github: https://github.com/vganusov/killing_in_gels.

## Ethics statement

No animal or human experiments performed.

## Author contributions

VVG and BM developed alternative models presented in the paper. The experimental data were generated by SB. The analysis was done primarily by BM. BM and VVG prepared the first draft of the manuscript and all the authors contributed to the final version.

## Acknowledgments

We would like to thank Dr. Sam Silverstein for giving us the permission to use their unpublished data from previous experiments (Datasets 4&5). their This work was supported by the NIH grants (R01GM118553 and R01AI158963) to VVG.

## Supplemental Information

**Supplemental Figure S1:**
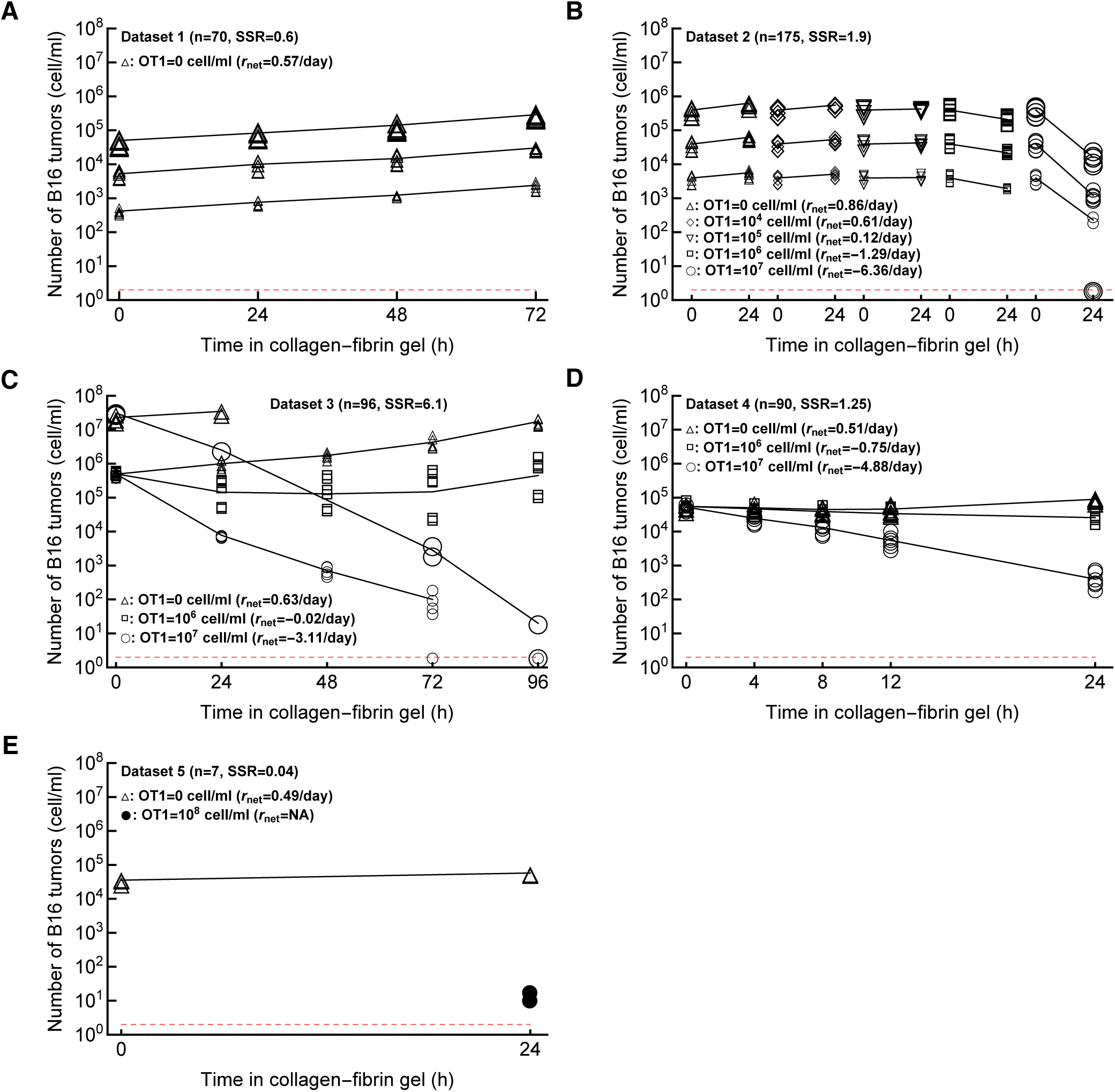
Data on the dynamics of B16 tumor cells for different time periods and at different CTL concentrations. We show all 5 datasets (Dataset 1-5, panels A-E) analyzed in this paper. (A) Dataset 1 (no CTLs) is on B16 tumor growth for 72 hours in the absence of CTLs; (B) Dataset 2 is on B16 tumor dynamics for 24 hours at different initial B16 cell and CTL concentrations (note that 5 gels had 0 B16 cells recovered, all at OT1= 10^7^ cells/ml); (C) Dataset 3 is on B16 tumor dynamics for up to 96 hours at different initial B16 cell and CTL concentrations (note that 8 gels had 0 B16 cells recovered at 72 and 96 hours post inoculation); (D) Dataset 4 on B16 tumor dynamics in the first 24 hours after inoculation at 3 different CTL concentrations, and (F) Dataset 5 (high CTL density) on B16 tumor dynamics for 24 hours at 0 and 10^8^ OT1 cells/ml. The size of markers indicates the different targeted number of B16 tumor cells. The lines connect average numbers (excluding gels with 0 B16 cells in B&C). For each panel we also show the number of gels *n* and sum of squared residuals (SSR) are computed by the relation 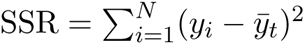. The red horizontal dashed line is the limit of detection for the experiments set at 2 cells/ml.

**Supplemental Figure S2:**
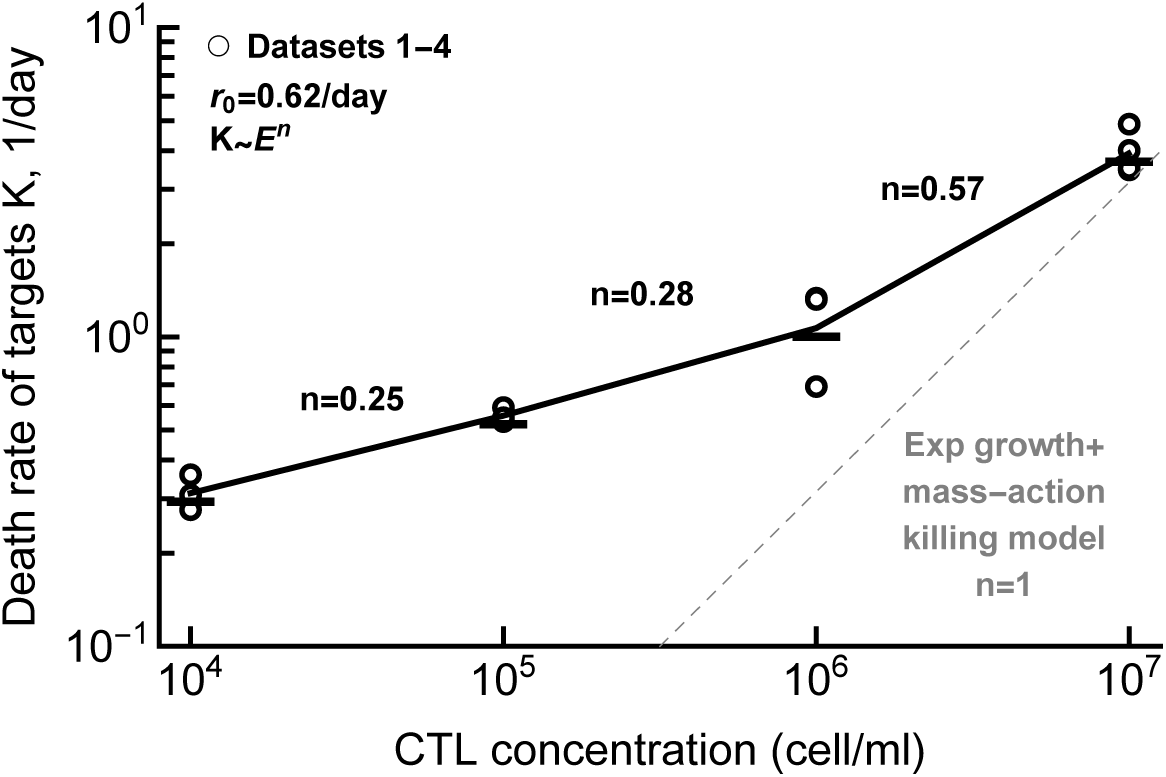
Regression analysis suggests nonlinear change of the death rate of B16 tumor cells with increasing CTL concentration. For the data in Datasets 1-4 we estimated the net rate of growth of B16 tumor cells over time *r*_net_ for every CTL and targeted B16 tumor concentrations (see Supplemental **Figure S1** for the average *r*_net_ per CTL concentration). In the absence of CTLs, the net growth rate of tumors was *r*_net_ = *r*_0_ = 0.62/day. We then calculated the death rate of B16 tumor cells *K* by substracting the estimated net rate of tumor change from *r*_0_, *K* = *r*_0_ *− r*_net_. Individual symbols are estimates of *K* for different target B16 tumor concentrations at a given CTL level. Assuming that death rate depends on CTL concentration as powerlaw with scale *n*, we estimated *n* for individual ranges of CTL concentrations. For example, the death rate of targets scales as *K ∼ E*^0.25^ for CTL concentrations *E* between 10^4^ and 10^5^ cells/ml. The dashed line shows a linear relationship *K ∼ E* between the death rate of targets *K* and CTL concentration *E* as predicted by the exponential-growth-mass-action-killing model (eqn. (3)).

**Supplemental Table S1:** The model with exponential growth of tumors and saturated killing rate by CTLs gives the best fit when the models are fitted to all data (Datasets 1-5). We list the best-fit parameters for the alternative models along with SSR, AIC, ΔAIC and Akaike weights *w*. Other details are similar to those given in Table 1.

| Datasets 1-5 ($E \leq 10^8$ cell/ml): n=438 | | | | | | | | | | | | |
| --- | --- | --- | --- | --- | --- | --- | --- | --- | --- | --- | --- | --- |
| Model | $\alpha$ | $r$, 1/day | $k$ | $h$ | $n$ | $g_0$ | $g_1$ | $g_2$ | SSR | AIC | ΔAIC | w |
| MA | 2.78 | 0.24 | $1.85 \times 10^{-7}$ | | | | | | 779 | 1503 | 779 | 0 |
| Sat | 2.81 | 0.696 | 7.32 | $8.63 \times 10^6$ | | | | | 131 | 724 | 0 | 1 |
| Power | 2.78 | 0.792 | 0.0017 |  | 0.477 |  |  |  | 147 | 776 | 52 | 0 |
| SiGMA | 2.95 | | $1.72 \times 10^{-7}$ | | | $2.88 \times 10^{-8}$ | 0.86 | 291 | 583 | 1380 | 656 | 0 |

**Supplemental Table S2:** Assuming different scaling factors *α* in best fit models moderately improves the fit but results in similar parameter estimates. We fitted the SiGMA model (eqn. (6)) to the data from Datasets 1-4 or the Sat model (eqn. (4)) to the data from Datasets 1-5 with one or five different scaling factors *α*.

| SiGMA model<br>Datasets 1-4 ($n = 431$) | | | Sat model<br>Datasets 1-4 ($n = 438$) | | |
| --- | --- | --- | --- | --- | --- |
| Parameters | Fixed $\alpha$ | Varied $\alpha$ | | Fixed $\alpha$ | Varied $\alpha$ |
| $\alpha$ | **2.71** | | | **2.82** | |
| $\alpha_1$ | | **3.18** | | | **2.89** |
| $\alpha_2$ | | **2.7** | | | **2.82** |
| $\alpha_3$ | | **2.74** | | | **2.86** |
| $\alpha_4$ | | **2.49** | | | **2.64** |
| $\alpha_5$ | | **3.85** | | | **3.56** |
| **r** |  |  |  | **0.7** | **0.7** |
| **k** | **$3.29 \times 10^{-7}$** | **$3.24 \times 10^{-7}$** |  | **7.2** | **7.2** |
| **h** | | | | $8.64 \times 10^6$ | **$8.14 \times 10^6$** |
| $g_0$ | **0.12** | **0.096** | | | |
| $g_1$ | **0.65** | **0.67** | | | |
| $g_2$ | **6714** | **6382** | | | |
| **AIC** | **654.2** | **650.5** |  | **723.7** | **727.5** |
| **LR** | **11.8** |  |  | **4.3** |  |
| $\chi(0.95,4)$ | **9.5** | | | **9.5** | |
| $p$ | **0.02** | | | **0.37** | |

**Supplemental Table S3:** A phenomenological Power model gives the best fit for the subset of the data. B16 tumor dynamics in two settings (at *T* = 10^6^ cell/ml and *E* = 10^6^ cell/ml from Dataset 3 and *T* = 10^5^ cell/ml and *E* = 10^6^ cell/ml from Dataset 4) is not monotonic (**Supplemental Figure S1**). We fitted 4 alternative models (eqns. (3)–(6)) to the subset of the data that excludes these two settings for Datasets 1-4 (top) or Datasets 1-5 (bottom). Other details are similar to those given in Table 1.

|  |  |  |  |  |  |  |  |  |  |  |  |  |
| --- | --- | --- | --- | --- | --- | --- | --- | --- | --- | --- | --- | --- |
| **Datasets 1-4** (subset) $n = 371$ | | | | | | | | | | | | |
| Models | $\alpha$ | **r** | **k** | **h** | **n** | $g_0$ | $g_1$ | $g_2$ | **SSR** | **AIC** | **ΔAIC** | **w** |
| MA | **2.88** | **0.72** | $3.84 \times 10^{-7}$ | | | | | | **88** | **526** | **99.7** | **0** |
| Sat | **2.74** | **0.72** | **4.8** | $2.49 \times 10^6$ | | | | | **71** | **451** | **24.7** | $10^{-6}$ |
| Power | **2.67** | **0.74** | **0.004** |  | **0.423** |  |  |  | **66.7** | **426.3** | **0** | **0.93** |
| SiGMA | **2.68** | | $3.17 \times 10^{-7}$ | | | $6.84 \times 10^{-8}$ | **0.72** | **7930** | **67.3** | **431.4** | **5.1** | **0.072** |
| **Datasets 1-5** (subset) $n = 378$ | | | | | | | | | | | | |
| Models | $\alpha$ | **r** | **k** | **h** | **n** | $g_0$ | $g_1$ | $g_2$ | **SSR** | **AIC** | **ΔAIC** | **w** |
| MA | **2.85** | **0.32** | $1.87 \times 10^{-7}$ | | | | | | **724** | **1327** | **889** | **0** |
| Sat | **2.86** | **0.72** | **9.36** | $1.39 \times 10^7$ | | | | | **82** | **503** | **65** | **0** |
| Power | **2.62** | **0.72** | **0.01** |  | **0.37** |  |  |  | **69** | **438** | **0** | **1** |
| SiGMA | **2.94** | | $1.76 \times 10^{-7}$ | | | $1.18 \times 10^{-7}$ | **0.84** | **252** | **544** | **1222** | **784** | **0** |

**Supplemental Table S4:** The Power model fits the subset of data best when we focus on a single targeted B16 tumor cell concentration in the gel. Here we divided Datasets 1-4 (top) or Datasets 1-5 (bottom) based on the target B16 concentration. For *T* = 10^4^ and 10^6^, the Power model provides the best fit. For *T* = 10^5^ without the high CTL data (Datasets 1-4), both the SiGMA and the Power model fits the data with similar Akaike weights. However, if we include the high CTL data (Datasets 1-5), the Sat model best explains the data. For other details of the table refer to Table 1.

|  |  |  |  |  |  |  |  |  |  |  |  |  |
| --- | --- | --- | --- | --- | --- | --- | --- | --- | --- | --- | --- | --- |
| Datasets 1-4 B16 = $10^4$ $n = 80$ | | | | | | | | | | | | |
| Models | $\alpha$ | r | k | h | n | $g_0$ | $g_1$ | $g_2$ | SSR | AIC | ΔAIC | w |
| MA | 2.74 | 0.6 | $3.55 \times 10^{-7}$ | | | | | | 14 | 96 | 60.5 | 0 |
| Sat | 2.67 | 0.62 | 4.08 | $1.85 \times 10^6$ | | | | | 8 | 54 | 18.5 | $10^{-4}$ |
| Power | 2.46 | 0.65 | 0.009 |  | 0.37 |  |  |  | 6.4 | 35.5 | 0 | 0.99 |
| SiGMA | 2.52 | | $2.93 \times 10^{-7}$ | | | $1.2 \times 10^{-7}$ | 0.67 | 8162 | 7.5 | 50 | 14.5 | $10^{-3}$ |
| Datasets 1-4 B16 = $10^5$ $n = 142$ | | | | | | | | | | | | |
| Models | $\alpha$ | r | k | h | n | $g_0$ | $g_1$ | $g_2$ | SSR | AIC | ΔAIC | w |
| MA | 2.42 | 0.53 | $4.63 \times 10^{-7}$ | | | | | | 20 | 134 | 36.65 | 0 |
| Sat | 2.36 | 0.58 | 6.48 | $4.07 \times 10^6$ | | | | | 16.5 | 107 | 9.65 | $4.2 \times 10^{-3}$ |
| Power | 2.33 | 0.58 | 0.001 |  | 0.52 |  |  |  | 15.37 | 97.35 | 0.17 | 0.48 |
| SiGMA | 2.34 | | $4.1 \times 10^{-7}$ | | | $1.37 \times 10^{-7}$ | 0.6 | 7322 | 15.14 | 97.18 | 0 | 0.52 |
| Datasets 1-5 B16 = $10^5$ $n = 149$ | | | | | | | | | | | | |
| Models | $\alpha$ | r | k | h | n | $g_0$ | $g_1$ | $g_2$ | SSR | AIC | ΔAIC | w |
| MA | 3.34 | 0.38 | $\times 10^{-7}$ | | | | | | 175 | 454 | 336.6 | 0 |
| Sat | 2.4 | 0.55 | 9.12 | $9.6 \times 10^6$ | | | | | 18 | 117.4 | 0 | 1 |
| Power | 2.35 | 0.62 | 0.02 |  | 0.33 |  |  |  | 22 | 149 | 31.6 | 0 |
| SiGMA | 2.96 | | $9.38 \times 10^{-8}$ | | | $1.38 \times 10^{-7}$ | 0.9 | 6106 | 139 | 425 | 307.6 | 0 |
| Datasets 1-5 B16 = $10^6$ $n = 112$ | | | | | | | | | | | | |
| Models | $\alpha$ | r | k | h | n | $g_0$ | $g_1$ | $g_2$ | SSR | AIC | ΔAIC | w |
| MA | 3.16 | 0.89 | $3.79 \times 10^{-7}$ | | | | | | 28 | 170 | 39 | 0 |
| Sat | 2.93 | 0.89 | 4.56 | $2.1 \times 10^6$ | | | | | 21.5 | 143 | 12 | $2 \times 10^{-3}$ |
| Power | 2.8 | 0.89 | 0.008 |  | 0.39 |  |  |  | 19.36 | 131 | 0 | 0.82 |
| SiGMA | 2.8 | | $2.95 \times 10^{-7}$ | | | $9.8 \times 10^{-8}$ | 0.9 | 10243 | 19.43 | 134 | 3 | 0.18 |

**Supplemental Figure S3:**
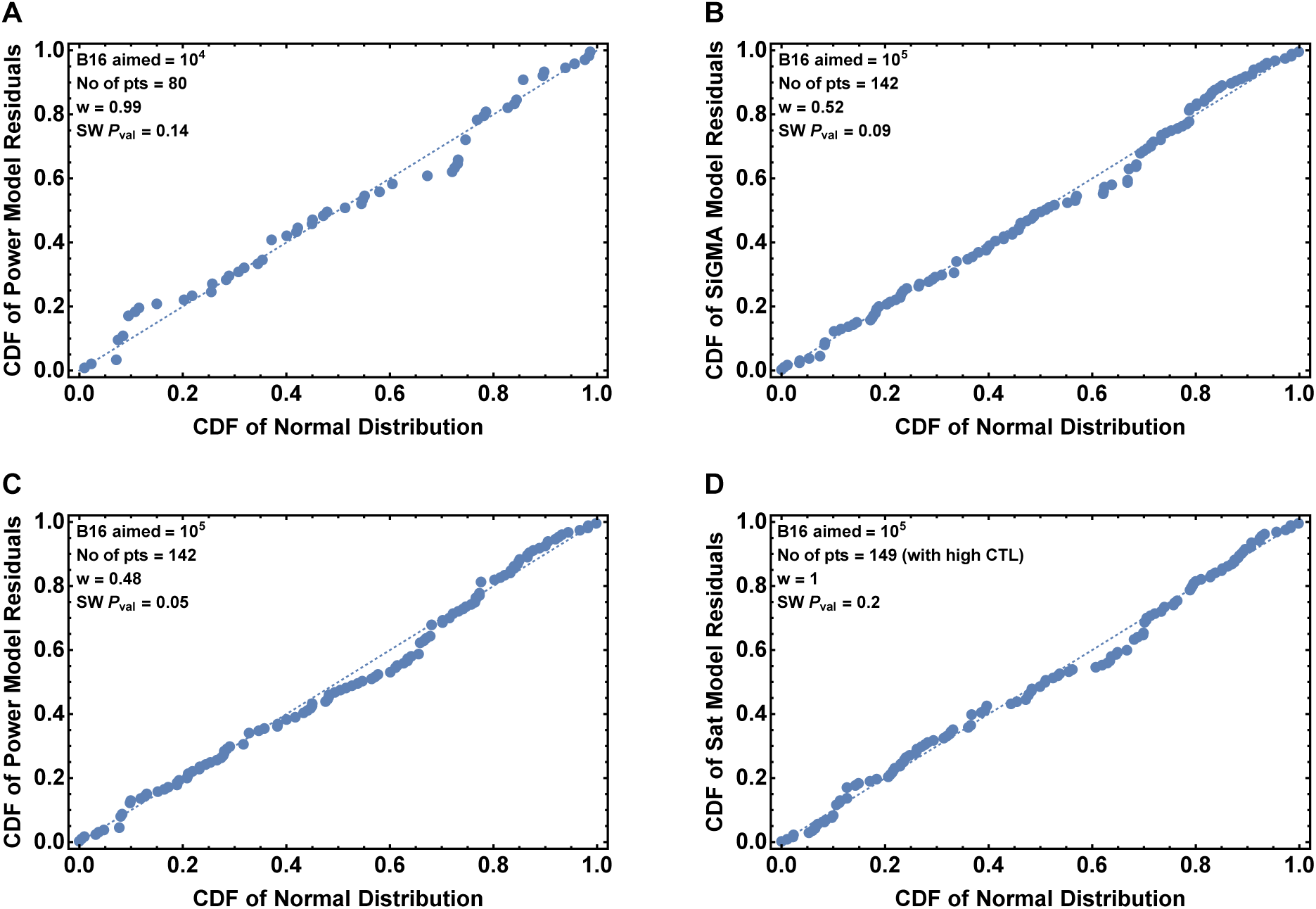
The residuals of the best models for sub-datasets with *T* = 10^4^ and 10^5^ are normally distributed. Here we show the normal probability plot of the best models of Table S4 for *T* = 10^4^ (A) and 10^5^ (B,C,D) with the p-value of the Shapiro-Wilk (SW) test.

**Supplemental Table S5:** The Power and the SiGMA models give the best fit if we fit the models to subsets of data experiment-wise. As we described in Materials and methods, each Datasets 1-4 has three experiments performed in duplicates. If we divide the data based on the three Experiments 1, 2 and 3 then the Power model gives the best fit for Experiment 1 and 3. For Experiment 2, the SiGMA model gives the best fit. The description of the table remain same as that of Table 1.

|  |  |  |  |  |  |  |  |  |  |  |  |  |
| --- | --- | --- | --- | --- | --- | --- | --- | --- | --- | --- | --- | --- |
| **Experiment 1** dataset $n = 125$ | | | | | | | | | | | | |
| Models | $\alpha$ | r | k | h | n | $g_0$ | $g_1$ | $g_2$ | SSR | AIC | $\Delta AIC$ | w |
| MA | 2.57 | 0.65 | $3.84 \times 10^{-7}$ | | | | | | 34 | 201 | 27 | 0 |
| Sat | 2.44 | 0.67 | 4.75 | $2.26 \times 10^6$ | | | | | 27.8 | 177 | 3 | 0.18 |
| Power | 2.39 | 0.67 | 0.003 |  | 0.44 |  |  |  | 27.15 | 174 | 0 | 0.8 |
| SiGMA | 2.43 | | $3.22 \times 10^{-7}$ | | | $9.6 \times 10^{-8}$ | 0.7 | 12726 | 28.4 | 182 | 8 | 0.015 |
| **Experiment 2** dataset $n = 126$ | | | | | | | | | | | | |
| Models | $\alpha$ | r | k | h | n | $g_0$ | $g_1$ | $g_2$ | SSR | AIC | $\Delta AIC$ | w |
| MA | 3.47 | 0.84 | $3.84 \times 10^{-7}$ | | | | | | 32 | 191 | 29.2 | 0 |
| Sat | 3.32 | 0.86 | 4.8 | $2.78 \times 10^6$ | | | | | 27 | 174 | 12.2 | 0.002 |
| Power | 3.2 | 0.86 | 0.005 |  | 0.42 |  |  |  | 25 | 164 | 2.2 | 0.25 |
| SiGMA | 3.18 | | $3.07 \times 10^{-7}$ | | | 0.018 | 0.84 | 6448 | 24.2 | 161.8 | 0 | 0.75 |
| **Experiment 3** dataset $n = 120$ | | | | | | | | | | | | |
| Models | $\alpha$ | r | k | h | n | $g_0$ | $g_1$ | $g_2$ | SSR | AIC | $\Delta AIC$ | w |
| MA | 2.69 | 0.67 | $3.84 \times 10^{-7}$ | | | | | | 18 | 121 | 61.6 | 0 |
| Sat | 2.55 | 0.7 | 4.8 | $2.45 \times 10^6$ | | | | | 12.5 | 79.5 | 20.1 | 0 |
| Power | 2.47 | 0.7 | 0.005 |  | 0.41 |  |  |  | 10.6 | 59.4 | 0 | 0.86 |
| SiGMA | 2.50 | | $3.22 \times 10^{-7}$ | | | $1.08 \times 10^{-7}$ | 0.72 | 8650 | 10.76 | 63 | 3.6 | 0.14 |

**Supplemental Figure S4:**
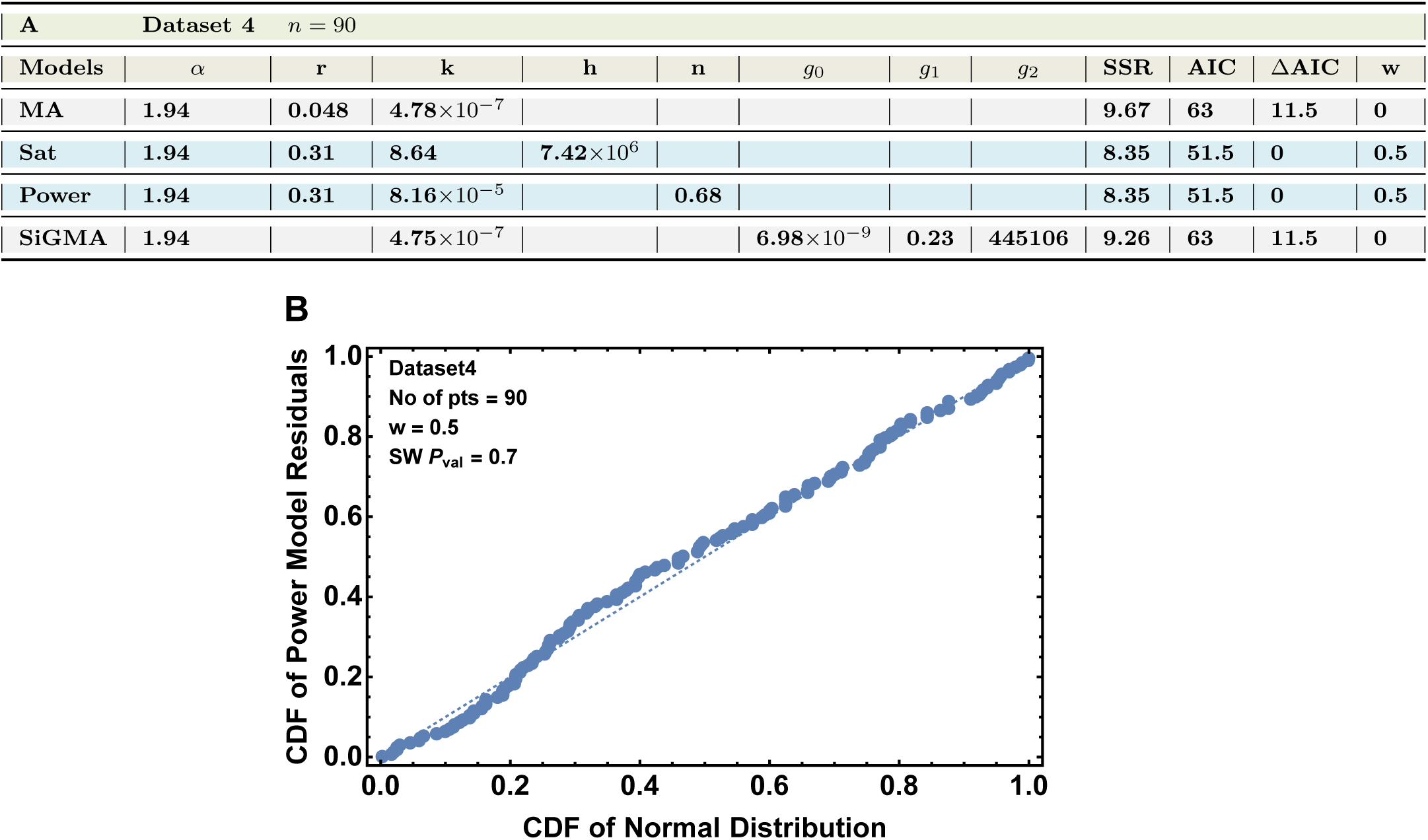
The phenomenological Power and the Sat models equally well describe the data for Dataset 4. Dataset 4 describes dynamics of B16 tumor cells within first 24 hours after inoculation into collagen-fibrin gels and has *n* = 90 data points. Parameter estimates are shown in panel A, and q-q plot for the the residuals for the models is shown in panel B. The table details in (A) are similar to Table 1.

**Supplemental Table S6:**
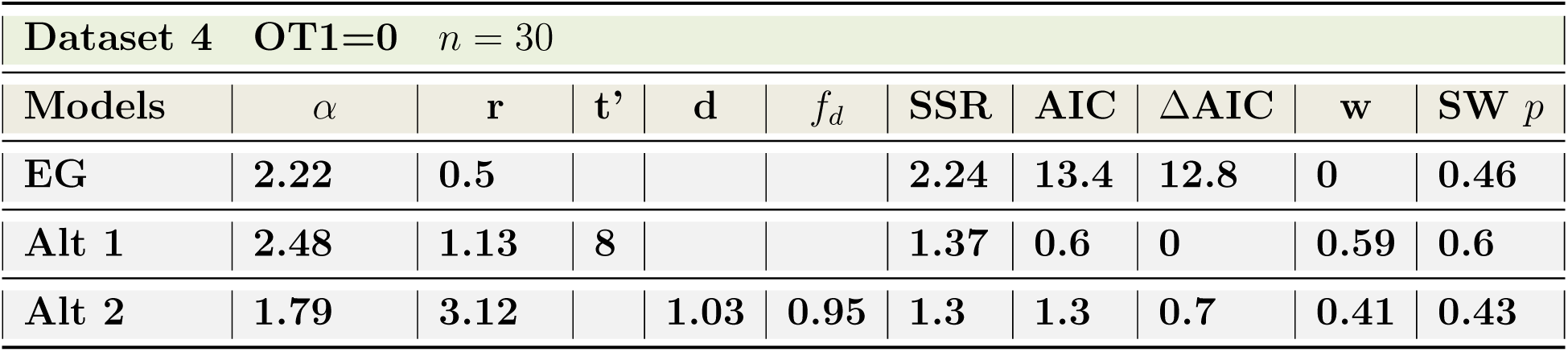
Both the alternative models fit the data better than the EG model for the growth only subset of the data in the Dataset 4. We selected the data on B16 tumor growth with OT1=0 resulting in *n* = 30 data points and fitted the EG, Alt 1, and Alt 2 models (eqn. (3) and eqns. (8)–(9), respectively) to these data (see **Figure 4B** for model fits). We show the results of the Shapiro-Wilk (SW) normality test of the residuals. Other details are similar to those in Table 1.

| Dataset 4 OT1=0 $n = 30$ | | | | | | | | | | |
| --- | --- | --- | --- | --- | --- | --- | --- | --- | --- | --- |
| Models | $\alpha$ | $r$ | $t'$ | $d$ | $f_d$ | SSR | AIC | ΔAIC | w | SW $p$ |
| EG | 2.22 | 0.5 |  |  |  | 2.24 | 13.4 | 12.8 | 0 | 0.46 |
| Alt 1 | 2.48 | 1.13 | 8 |  |  | 1.37 | 0.6 | 0 | 0.59 | 0.6 |
| Alt 2 | 1.79 | 3.12 |  | 1.03 | 0.95 | 1.3 | 1.3 | 0.7 | 0.41 | 0.43 |

**Supplemental Figure S5:**
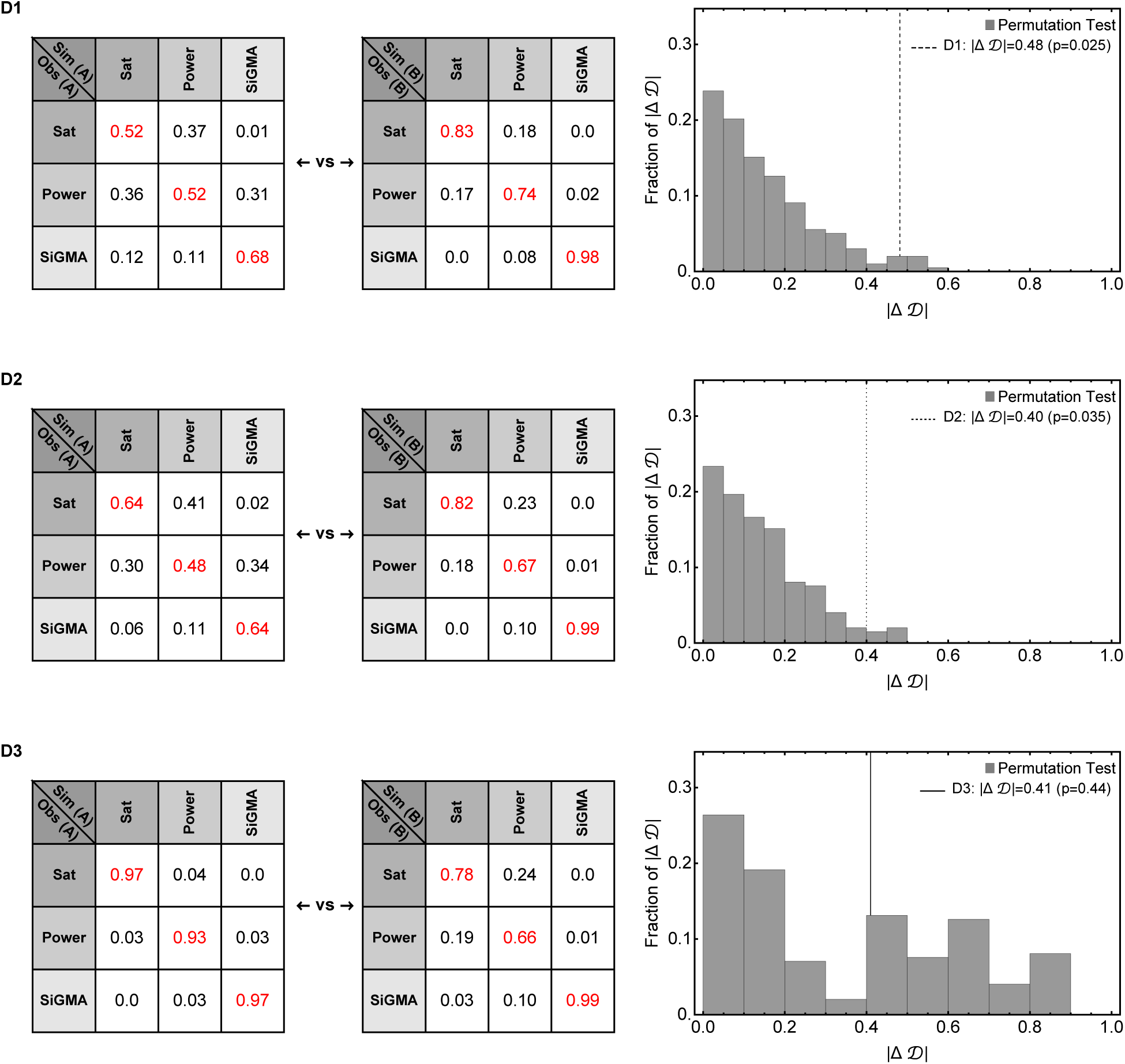
Statistical power to detect a difference in the fit quality between alternative mathematical models depends on experimental design. We performed simulations of 3 experimental designs measuring impact of CTLs on B16 tumor dynamics (see **Figure 5** and Main text for details). For designs D1 and D2 we show that the experiment type A and B are significantly different from each other. With permutation test, however, for D3 we fail to reject the null hypothesis that the experiments are similar. For three simulated experimental designs D1, D3 and D3 we simulated 100 identical replicas for investigation Type A and B from a model while choosing the errors randomly and then fitted them with models. This allowed us to get matrices like the ones in the left 2 panels. The red diagonal entries show fraction of replicas generated by the a model is also best fitted by the same model where as the off diagonal entries present fraction of replicas generated by a model but best fitted by a different model. The experimental Type A or B with heavier diagonal terms would indicate a better experiment. In this plot we did a permutation test to compare the observed |Δ*Δ|*_obs_ in a permutated distribution of |Δ*Δ|*_per_ to obtain a p-value, where *Δ* is a determinant of the matrices. This test allowed us to statistically comment on the structural difference of the design Types A and B. The details of the test is discussed in the end of Results section. See eqn. (12) for test statistic measure.

## Notes

https://github.com/vganusov/killing_in_gels

### Competing Interest Statement

The authors have declared no competing interest.

